# The Rice *propiconazole resistant 1-D* mutant, with activated expression of a DPb transcription factor gene, exhibits increased seed yields

**DOI:** 10.1101/2021.01.02.425087

**Authors:** Claudia Corvalán, Gynheung An, Sunghwa Choe

## Abstract

Mutants defective in brassinosteroid (BR) biosynthesis or signaling pathways often display semi-dwarfism, as do the highly productive gibberellin mutants that enabled the Green Revolution. However, reduced vegetative growth in BR mutants does not necessarily correspond to increased seed yields. To better understand the mode of action of BR, we isolated a rice *propiconazole resistant1-D* (*pzr1-D*) mutant by screening an activation-tagging mutant population in the presence of the BR biosynthesis inhibitor propiconazole (Pcz). The expression of a putative transcription factor gene homologous to Arabidopsis Dimerization Partner (*DPb*) was activated in *pzr1-D. pzr1-D* exhibited characteristic phenotypes such as reduced height, and increased seed yields and tiller numbers. Like Arabidopsis *DPb*, rice *PZR1* is expressed differentially in the tissues examined. Furthermore, *pzr1-D* displayed altered cell division phenotypes, including the production of small calli. In addition, the cell number and size in mutant roots and leaves differed from those in wild-type plants of the same age. RNA sequencing revealed that the promoters of differentially expressed genes are enriched with cognate sequences for both BZR1 and EF-DPb transcription factors, suggesting that PZR1 functions in BR-mediated cell division in rice. *PZR1* expression may thus be manipulated to increase seed yield in economically important rice varieties.

**SIGNIFICANCE STATEMENT:** Mutants defective in brassinosteroid biosynthesis or signaling pathways often display semi-dwarfism. However, in many cases these mutants do not necessarily produce increased seed yields. We show a rice *propiconazole resistant1-D* (*pzr1-D*) mutant which exhibits reduced height phenotypes along with increase tiller number and seed yields. *pzr1-D* shows activation of a putative homologous to Arabidopsis Dimerization Partner (*DPb*) with functions in cell division, pointing that PZR1 may function in BR-mediated cell division in rice.

## INTRODUCTION

Rice serves as both a staple food and a model plant for molecular studies. Many studies have focused on improving traits that are associated with increased seed yield, including plant height, leaf erectness, number of tillers and panicles, and seed size. Significant progress has recently been made through the analysis of activation-tagging or knockout mutants, including studies showing that brassinosteroids (BRs) play crucial roles in controlling plant architecture (Clouse *et* al., 1996, Choe *et al*., 1998, Yamamuro *et al*., 2000, Sakamoto *et al*., 2006).

Typical BR-deficient mutants in rice display dwarf phenotypes, including dark-green, erect leaves and shortened leaf sheaths in the early vegetative stage of growth. After flowering, the mutant plants are only ∼40% the height of wild-type plants, and internode elongation, especially the second internode, differs from that of the wild type, with malformed panicles and a reduced number of branches and spikelets (Hong *et al*., 2003, Tanabe *et al*., 2005, Nakamura *et* al., 2006). By contrast, plants overexpressing BR biosynthesis genes or plants with increased BR sensitivity often have a large stature, with increased numbers of flowers and seeds and lamina with increased bending from the vertical axis of the leaf towards the abaxial side (Wu *et al*., 2008, Tanaka *et al*., 2009, Zhang *et al*., 2009). Regulating the expression of genes involved in modulating endogenous BR levels or affecting sensitivity to BR is a promising technique for improving agricultural traits. To date, the expression of the key BR biosynthetic gene, *DWF4*, has been altered in *Arabidopsis thaliana, Oryza sativa* (rice), tomato, *Brassica napus* and maize (Choe *et al*., 2001, Sakamoto *et al*., 2006, Liu *et al*., 2007, Li *et al*., 2016, Sahni *et al*., 2016). Similarly, studies in the BR receptor BRASSINOSTEROID INSENSITIVE 1 (BRI1) mutant *d61* in rice and *zmbri1*-RNAi plants in maize has been reported (Morinaka *et al*., 2006, Kir *et al*., 2015). In these examples, the overexpression or disruption of *DWF4* or *BRI1* resulted in plants with desirable traits like increased seed yield. Thus, identifying novel mutants related to BR biosynthesis or action could reveal novel ways to enhance seed yield phenotype.

Brassinazole (Brz) is a BR biosynthesis inhibitor that has been used to help identify novel components of the BR biosynthesis and signaling pathways in Arabidopsis (Wang *et al*., 2002, Kim *et al*., 2014, Maharjan *et al*., 2014). Similarly, we recently employed BR sensitivity screening in rice using the triazole-type inhibitor propiconazole (Pcz) (Corvalan and Choe, 2017). In contrast to costly Brz, Pcz is a commercially-used fungicide that is readily accessible and inexpensive, allowing it to be used in large-scale chemical genomics and field-testing. Pcz treatment produces typical BR-deficient phenotypes, such as epinastically growing, dark-green cotyledons, and reduced growth of hypocotyls and primary roots (Hartwig *et al*., 2012). Pcz treatment of genetically disrupted *dwf4-1* mutant seedlings suggested that DWF4 is likely a target of Pcz (Asami *et al*., 2001, Chung *et al*., 2011, Hartwig *et al*., 2012).

Plants respond to environmental cues by altering their growth. Because plant growth depends on both cell elongation and division, the regulation of the cell cycle is particularly important. The progression of the cell cycle has two major checkpoints: the transition from G_1_ to S phase and from G_2_ to M phase. The E2F family of transcription factors regulates the transcription of genes involved in the G_1_-to-S phase transition. The DNA-binding activity of E2F is stimulated by binding to Dimerization Partner (DP) proteins; E2F-DP heterodimeric transcription factors activate the expression of genes responsible for cell cycle control, the initiation of replication, and enzymes required for DNA synthesis during S phase (Kosugi and Ohashi, 2002). The E2F-DP pathway is conserved in animals and plants. In Arabidopsis, at least three E2Fs (E2Fa, E2Fb, and E2Fc) and two DPs (DPa and DPb) have been identified, along with their target genes (Magyar *et al*., 2000, Kosugi and Ohashi, 2002, Vandepoele *et al*., 2005). The overexpression of *E2F* and *DPa* in Arabidopsis produced dwarfed plants with curled leaves and cotyledons (De Veylder *et al*., 2002). A genome-wide analysis identified several core cell cycle genes in rice, including four E2Fs and three DPs (*OsDP1, OsDP2*, and *OsDP3*). However, only *DP1* transcripts have been detected by RT-PCR, while attempts to detect *DP2* and *DP3* transcripts were not successful (Guo *et al*., 2007).

In the current study, we screened a T-DNA activation-tagging mutant population in the presence of Pcz treatment for Pcz-resistant lines. These mutants were developed using the pGA2715 vector harboring four copies of the constitutive CaMV 35S enhancers, which can cause transcriptional activation of genes flanking the inserted T-DNA, resulting in dominant gain-of-function mutations (Jeong *et al*., 2002). Among the 17 propiconazole-resistant lines recovered, we further analyzed the *propiconazole-resistant 1* (*pzr1-D*) mutant with the phenotypes of increased seed yields. This mutant exhibits several characteristic BR phenotypes, such as semi-dwarfism and a greater number of tillers along with increased sensitivity to BRs. Molecular characterization and phylogenetic analysis showed that the inserted T-DNA in *pzr1-D* activates the expression of a homolog of the Arabidopsis *DPb* transcription factor gene involved in cell cycle regulation. Both dominant mutants and transgenic lines overexpressing *PZR1* showed increased numbers of tillers, panicles, and branches in panicles, all contributing to a considerable increase in seed yield. Our findings reveal a possible role for PZR1 in mediating BR-regulated cell division control in rice. These results may serve as a basis for developing novel approaches for increasing seed yield.

## RESULTS

### Mutation by activation tagging confers resistance to the BR biosynthesis inhibitor propiconazole

We previously reported that Pcz is a potent, specific BR inhibitor and demonstrated its use in maize and *Brachypodium* (Hartwig *et al*., 2012, *Corvalan and Choe, 2017)*. In contrast to its costly counterpart, Brz, the low cost of Pcz allowed us to perform large-scale tests to screen a T-DNA activation-tagging mutant population. These mutants were developed using the T-DNA vector pGA2715 with 35S enhancers, which can cause transcriptional activation of genes flanking the insertion, resulting in dominant gain-of-function mutations (Jeong *et al*., 2002).

First, we investigated the response of rice to various concentrations of the inhibitor to identify the optimal conditions for isolating Pcz-resistant mutants. As expected, Pcz affected plant growth and induced dwarfism in a dose-dependent manner (Figure S1A and S1B). Treatment with 30 μM Pcz reduced the total size of wild-type seedlings by up to 47%, and the response of roots was even more severe, with a 65% inhibition of growth (Figure 1A and 1B). To ensure that Pcz was effectively inhibiting BRs, we examined the expression levels of a key BR biosynthetic gene, *OsDWF4. OsDWF4* mRNA levels increased in plants treated with Pcz, which is in accordance with the negative feedback regulation of BR biosynthetic genes (Figure 1C). Thus, we expected that under this treatment, we could perform visible phenotypic screening for resistant seedlings, as reflected by their longer roots and/or leaves compared to wild-type seedlings.

**Figure 1.**
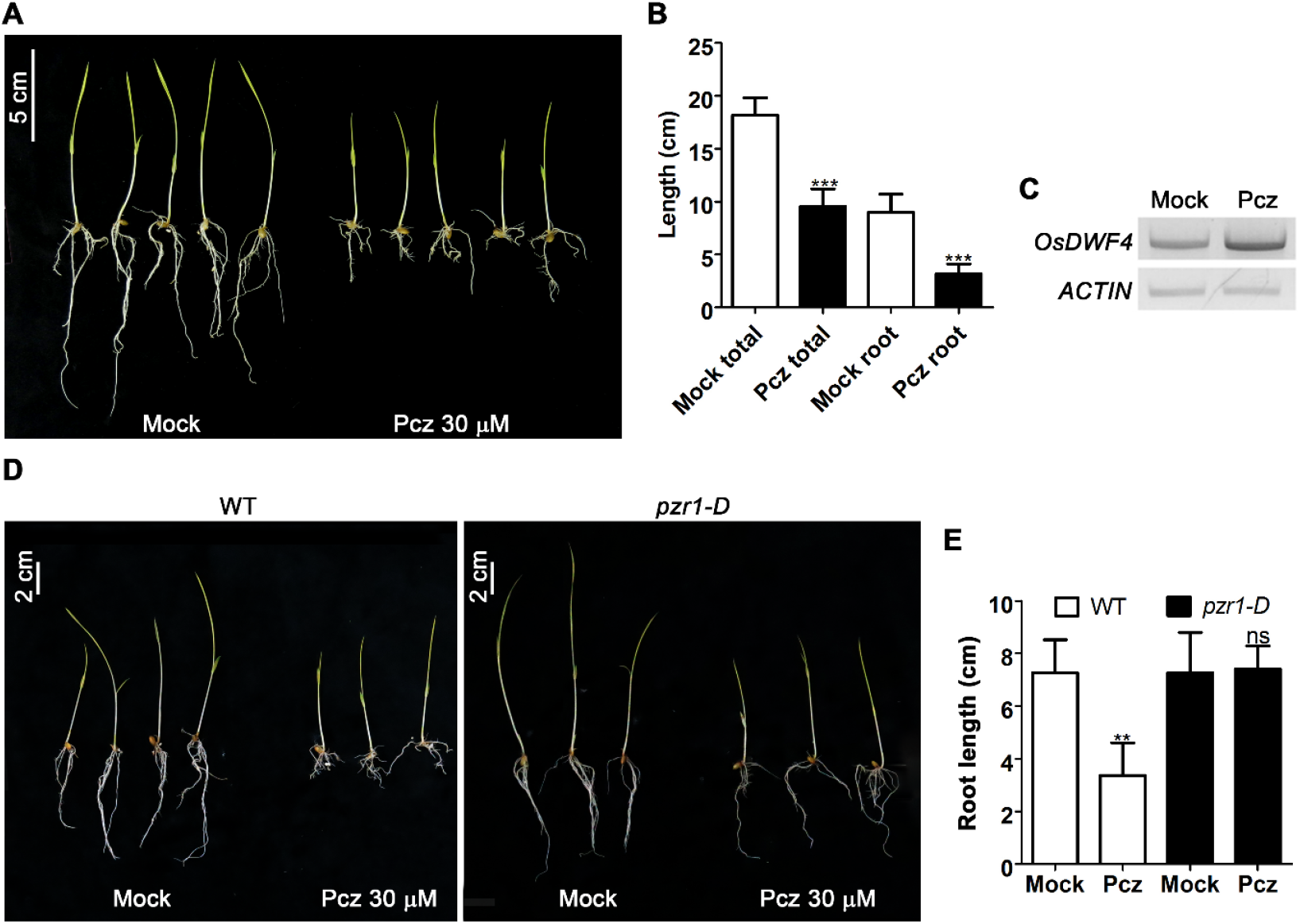
Propiconazole resistance of *pzr1-*D mutant seedlings. (**A**) Morphology of 10-day-old rice seedlings under mock or 30 μM Pcz treatment in darkness. (**B**) Total lengths and root lengths of seedlings shown in panel A. Values represent the average of at least 10 samples, and error bars represent standard deviation. (**C**) RT-PCR comparing *DWF4* expression in mock and Pcz treatments. Actin was used as an internal control. (**D**) Morphology of 10-day-old wild-type and mutant rice seedlings under mock or 30 μM Pcz treatment. (**E**) Root lengths of wild-type and *pzr1-D* seedlings under each treatment. Values represent the mean of 12 samples per treatment. Error bars represent standard deviation. In both graphs, significant differences among treatments were determined using Student’s *t*-test. **, P<0.001; ***, P<0.0001; and ns, non-significant.

We identified 17 lines from the activating-tagging mutant population with various degrees of resistance to the treatment, which we designated PROPICONAZOLE RESISTANT (PZR) 1 to 17. Among these lines, we isolated a dominant mutant, *pzr1-D*, with visible resistant phenotypes that co-segregated with the T-DNA insertion (Figure S2A). The root lengths of homozygous mutant seedlings under 30 μM Pcz treatment were almost identical to those under mock conditions, unlike the wild type, which exhibited a 45% inhibition of root growth (Figure 1D and 1E). Pcz treatment under dark conditions yielded similar resistant phenotype as to those obtained under the light (Figure S1C and S1D).

### *PZR1* regulates plant architecture and yield in rice

We examined the phenotypes of adult plants in the field and under greenhouse conditions and found that the *pzr1-D* plants had higher yields than wild-type plants (Figure 2). The total plant weight was 33% higher in the mutant than in the wild type, although their total heights did not significantly differ. Increases in weight can be explained by the increased number of tillers in the mutant, as well as the increased number of panicles per plant (Figure 2A to 2E). Seed weight per plant increased from 30 g in wild type to 50 g in the mutant, indicating an increase in yield of approximately 160%. We then examined the morphology of the panicles and found that not only was the number of panicles increased in the mutant, but the number of primary and secondary branches per panicle was also higher in *pzr1-D* than in the wild type (Figure 2F to 2I). We reasoned that the higher weight of total seeds in the activation-tagging mutants is likely due to an increased number of seeds per plant rather than an increase in seed size. Indeed, the seed size, area, and weight of mutant seeds were not significantly different from those of wild-type seeds. In fact, the mutant seeds were slightly smaller than wild-type seeds (Figure S3).

**Figure 2.**
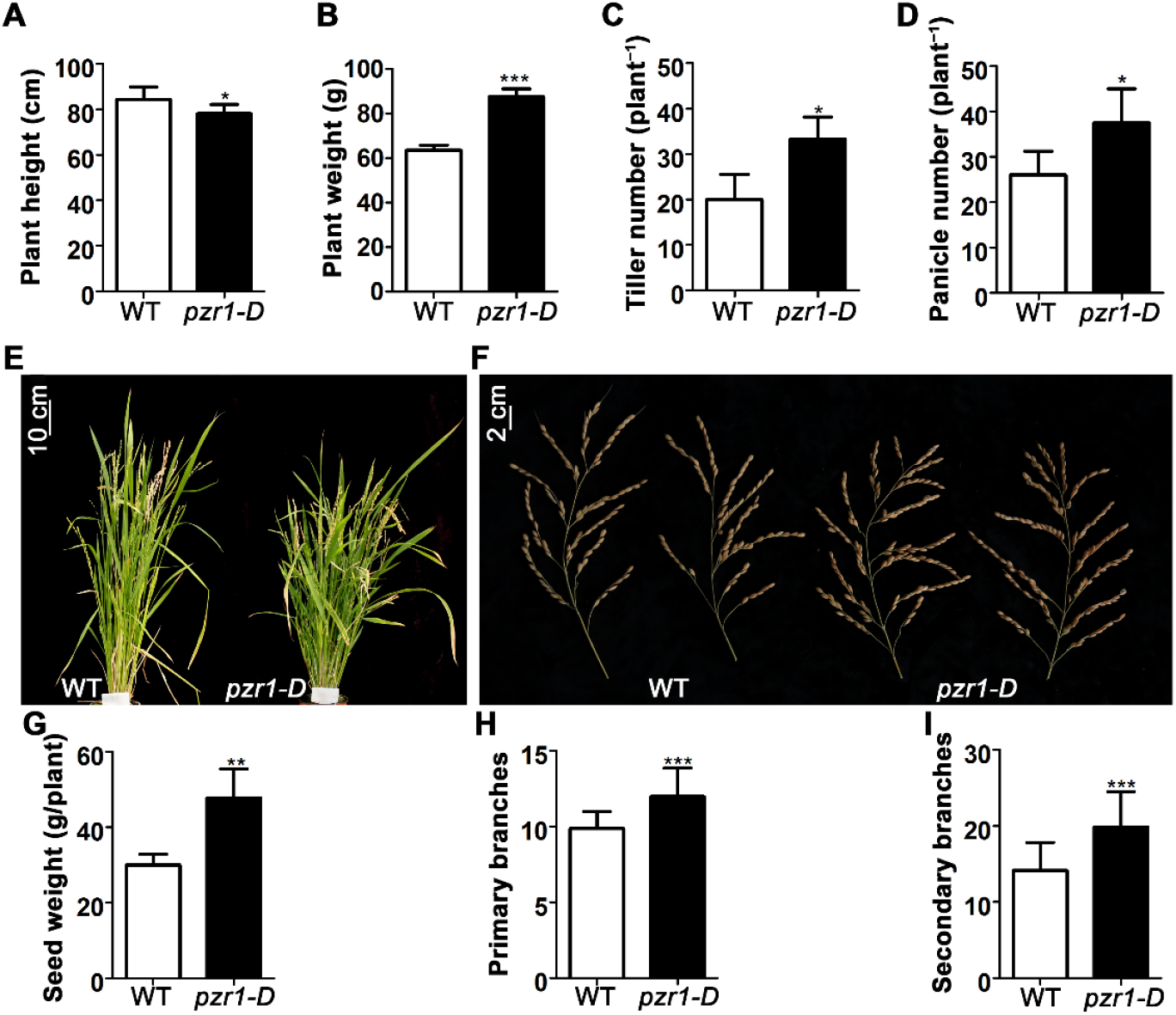
Phenotypes of *pzr1-D* adult plants. (**A**) Plant height, (**B**) plant weight, (**C**) tiller number, and (**D**) number of panicles was plotted for wild type and *pzr1-D*. (**E**) Comparison of adult plants and (**F**) panicle morphology between the wild type and mutant. Comparison of seed weight per plant, (**H**) number of primary branches, and (**I**) number of secondary branches. The graphs represent average values (n>7), and error bars represent standard deviation among samples. Significant differences among treatments were determined using Student’s *t*-test. *, P<0.05; **, P<0.001; and ***, P<0.0001.

### Mutant plants show increased sensitivity to BR

Since the *pzr1-D* mutant was isolated based on its resistance to a BR inhibitor, we evaluated the response of the mutant to exogenous brassinolide (BL) treatment. Rice leaf bending is sensitive to active BRs, which forms the basis for the well-known lamina-joint inclination bioassay to investigate BR responses (Wada *et al*., 1981). Under conventional growth conditions, the bending angle of the leaves of *pzr1-D* seedlings was greater than that of wild-type plants (Figure 3A and 3B). In the lamina bending assay, treatment with BL led to a dramatically increased leaf angle in *pzr1-D* plants, whereas wild-type plants exhibited a milder response (Figure 3C and 3D). At greater BL concentrations, the difference in the leaf angle response became more pronounced (Figure 3E). These results suggest that the *pzr1-D* mutant is more sensitive to exogenous BL treatment than are wild-type seedlings. To confirm this notion, we investigated root and coleoptile growth of seedlings in response to BL treatment under dark conditions. In the absence of BL, there were significant differences in root growth between *pzr1-D* and its wild-type counterpart when grown in darkness (Figure 3F). Moreover, when the medium was supplemented with BL, the inhibition of root growth was more pronounced in *pzr1-D* than in the wild type, and the opposite response was observed in coleoptiles (i.e., increased growth) (Figure 3G and 3H). Thus, the increased sensitivity of the mutant seedlings was confirmed based on their reduced root growth and increased coleoptile elongation in response to BL.

**Figure 3.**
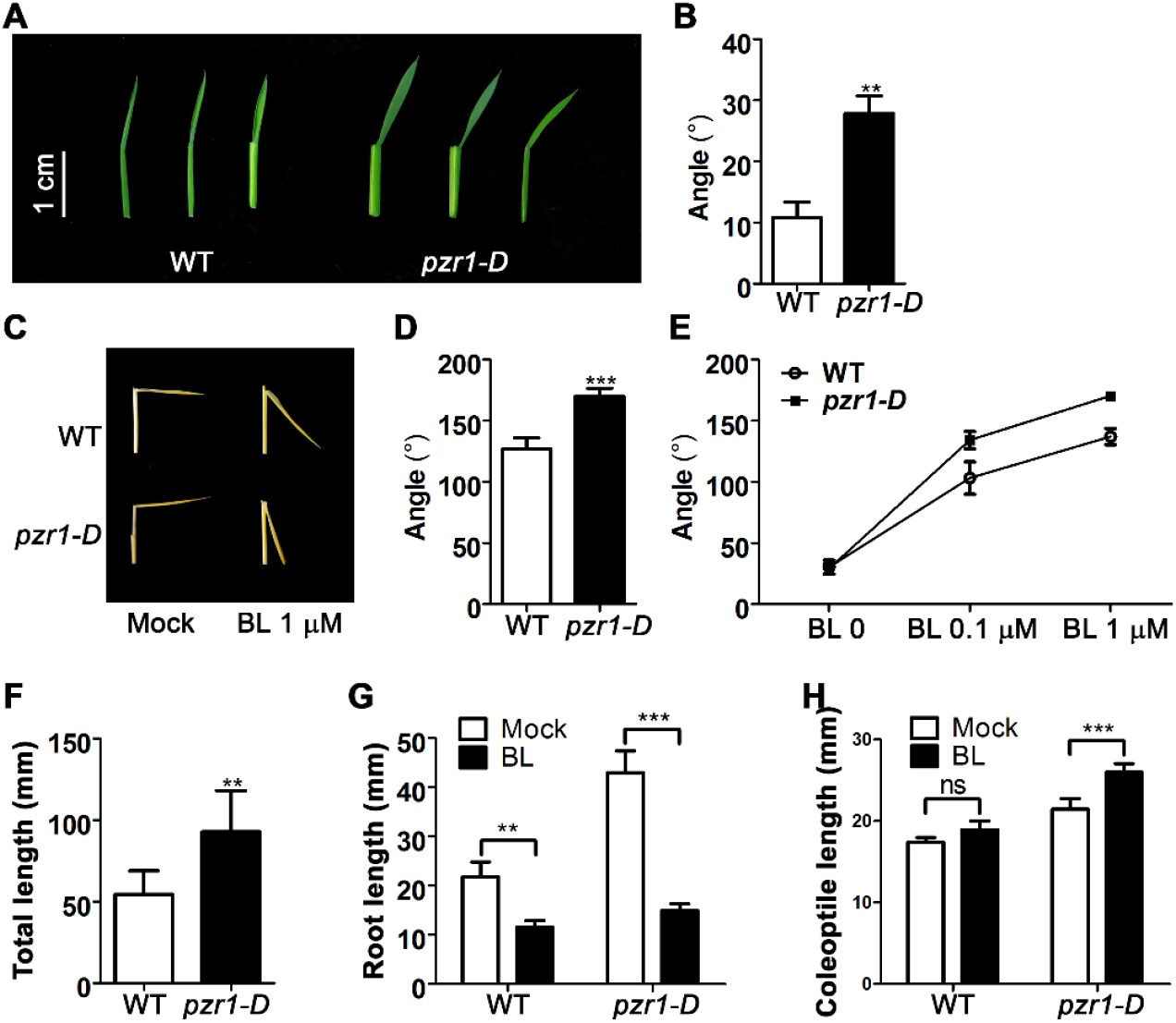
BR-related phenotypes of *pzr1-D* mutant seedlings. (**A–B**) Inclination of the segment corresponding to the second leaf from wild-type and *pzr1-D* mutant plants. (**C–E**) Lamina inclination bioassay performed using the indicated concentrations of brassinolide (BL). The graphs represent average values (n=15), and error bars represent standard deviation among samples. (**F**) Effect of darkness on wild-type and *pzr1-D* seedlings. (**G**) BL sensitivity tested by root inhibition in various genotypes in the presence of BL (1 μM) and in darkness. Differences in coleoptile growth in seedlings in response to BL and dark treatment. Significant differences among treatments were determined using Student’s *t*-test. **, P<0.001; ***, P<0.0001; and ns, non-significant.

### The *pzr1-D* mutant shows altered cell number and size in different tissues

We observed root and leaf tissues of wild-type and mutant seedlings under a confocal microscope and noticed abnormalities in the *pzr1-D* samples. Roots of *pzr1-D* contained greater number of cells, most of which were smaller than those in wild-type roots (Figure 4A to 4D). Similarly, mutant leaves were wider relative to wild-type leaves and had more but smaller cells (Figure S4). Because cell division appeared to be altered in the mutant, we examined callus initiation and morphology in wild-type and *pzr1-D* plants. Callus induced from mutant seeds was smaller than that derived from wild-type seeds, and it exhibited adventitious root formation, whereas wild-type callus did not (Figure 4E and 4F). Therefore, *PZR1* might play a role in regulating the cell division in rice.

**Figure 4.**
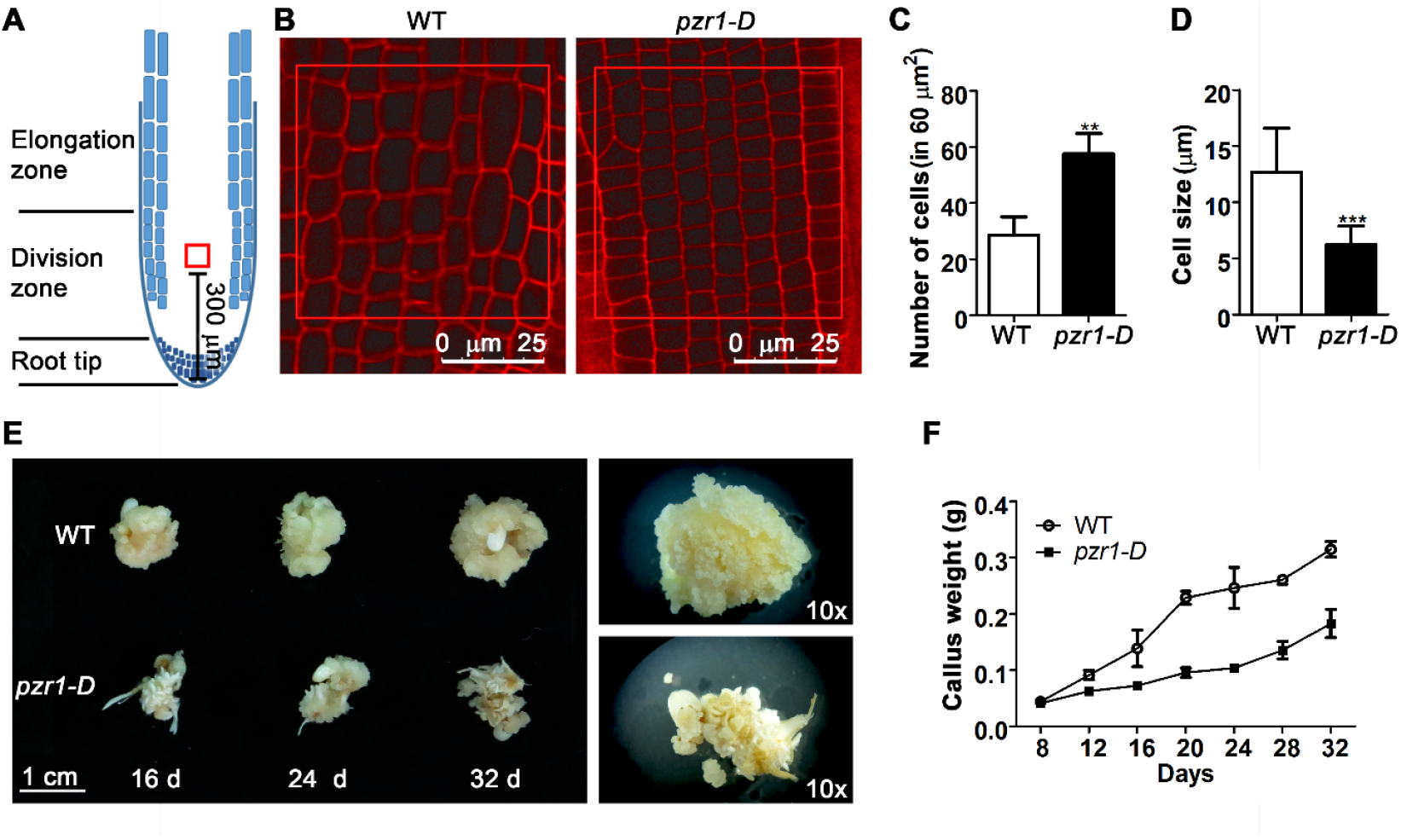
Microscopy analysis of *pzr1-D* and morphology of calli derived from mutant and wild-type seeds. (**A**) Schematic representation of a root. Red square (60 mm^2^) shows the point at which the images **(B)** were taken and the cells counted **(C)** and measured **(D)**. (**B**) Images show a section of the meristematic zone of PI-stained roots of 7-day-old wild-type and *pzr1-D* seedlings. (**C**) Number and (**D**) size of the cells contained in the square were counted in four different samples per genotype, and the average values with standard deviation were plotted. (**E**) Morphology of calli derived from wild-type and *pzr1-D* plants at the indicated time. (**F**) Growth profile showing the effects of the *pzr1-D* mutation on callus development. Error bars represent standard deviation, and significant differences were determined using Student’s *t*-test. *, P<0.05; **, P<0.001; and ***, P<0.0001.

### Activation tagging and overexpression of the rice homolog of Arabidopsis *DPb* underlie the phenotypes observed in *pzr1-D*

Previous analyses of the activation-tagging mutant population used in this study (Jeong *et al*., 2002) revealed that a gene, *Os03g05760*, is located 1.8 kb upstream of the T-DNA insertion (Figure 5A). We therefore investigated the expression level of *Os03g05760* and two other genes (*Os03g05750* and *Os03g05770*) near the insertion. The expression level of *Os03g05760* in the mutant was approximately 10-times that in the wild type, whereas the expression levels of the other two genes were like those of wild-type samples (Figure 5B). Thus, the increase in *Os03g05760* mRNA levels appears to be responsible for the mutant phenotypes in *pzr1-D*. We also examined the expression level of this gene in heterozygous and segregating wild-type plants and found that its expression increased in the mutant in a gene dose-dependent manner (Figure 5C). This, along with the intermediate phenotypes observed in heterozygous plants, indicates that the mutation is dominant: we therefore designated it as *pzr1-D* (Figure S2B to S2F).

**Figure 5.**
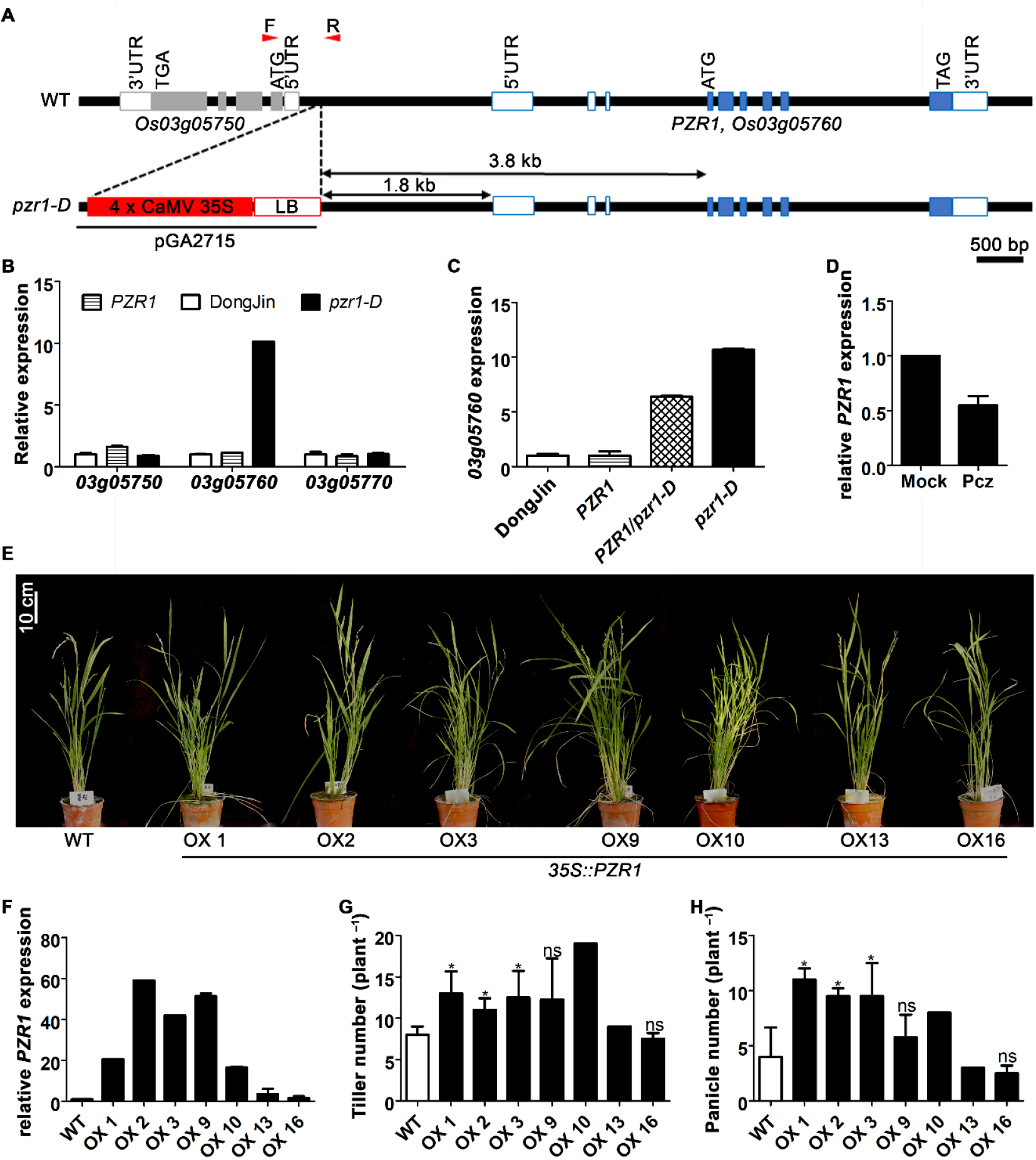
Activation-tagging T-DNA insertion in *PZR1* and *PZR1* overexpression rice lines. (**A**) Schematic representation of *PZR1* wild-type (WT) and mutant (*pzr1-D)* alleles. Arrows indicate the distance from the border of the insertion to the gene. White boxes represent 5’ and 3’ UTRs, blue boxes represent exons, red box define the 35S promoter in the T-DNA and black lines represent introns. Red arrowheads indicate the positions of forward and reverse primers used for genotyping. Gray boxes represent exons of the gene upstream of *PZR1*. (**B**) RT-qPCR of three genes adjacent to the T-DNA insertion. (**C**) Transcription levels of *PZR1* in the wild-type (DongJin), segregating wild-type (*PZR1*), heterozygous (PZR1/*pzr1-D*), and mutant (*pzr1-D*) genotypes. (**D**) RT-qPCR analysis of *PZR1* expression in the wild type (WT) after propiconazole (30 μM) treatment. Graphs show data from a representative experiment out of three biological replicates performed. (**E**) Morphology of representative 1-month-old plants from each line. (**F**) Number of tillers, (**G**) number of panicles, and (**H**) expression levels of *PZR1* in transgenic plants compared with non-transformed plants. Error bars represent standard deviation, and significant differences were determined using Student’s *t*-test. *, P<0.05; ns, non-significant.

We then investigated whether *PZR1* expression is affected by BL levels by measuring transcript levels in seedlings following Pcz treatment. *PZR1* expression was downregulated in plants subjected to this treatment (Figure 5D).

To confirm that the observed phenotypes were caused by the increased expression of *PZR1* in the *pzr1-D* mutant, we transformed wild-type rice and Arabidopsis plants with a vector expressing the *PZR1* coding sequence (CDS) under the control of the CaMV 35S promoter. In rice plants, like in *pzr1-D*, the tiller and panicle number increased in lines significantly overexpressing *PZR1* (OX 1–10), especially in OX 1, 2, 3, 9, and 10. The expression levels of this gene in lines #13 and #16 were not very different from those of the wild type, and their phenotypes were similar to those of Dongjin plants (Figure 5E to 5H. For Arabidopsis *PZR1*-overexpressing plants, seedlings with high expression levels of *PZR1* (i.e., OX 2, 3, and 5) showed increased lateral root number, which could be attributed to the role of *DP* in the cell cycle (Figure S5).

Next, we investigated the possible functions of *PZR1/Os03g05760*. We screened the TAIR Arabidopsis database using the full-length protein sequence of the product of *Os03g05760* (Q84VA0) for homologous genes. The highest scores corresponded to two DP (Q9FNY2 and Q9FNY3) and three E2F (Q9FV71, Q9FV70, and F4ILT1) proteins, all from a family of transcription factors with roles in the cell cycle. Plant DP proteins have been identified in Arabidopsis and *Triticum aestivum* (Magyar *et al*., 2000, Ramirez-Parra and Gutierrez, 2000), and three possible DP homologs in rice have been found using genome-wide analysis (Guo *et al*., 2007). Phylogenetic analysis of the PZR1 sequence, along with those of known DP proteins in plants, supported the idea that PZR1 is likely a homolog of Arabidopsis DPb (Figure 6A). Thus, to investigate the functions of *PZR1* in rice, we examined its expression pattern in various tissues from seedlings and adult rice plants (Figure 6B to 5E). The expression level of this gene was relatively low in seedlings, with lower expression levels in roots compared to aerial tissues (Figure 6B and 6C). Nonetheless, we detected these transcripts in all adult plant tissues examined, with leaf tissues having the highest expression levels. This expression pattern was even more pronounced in the *pzr1-D* mutant (Figure 6D and 6E).

**Figure 6.**
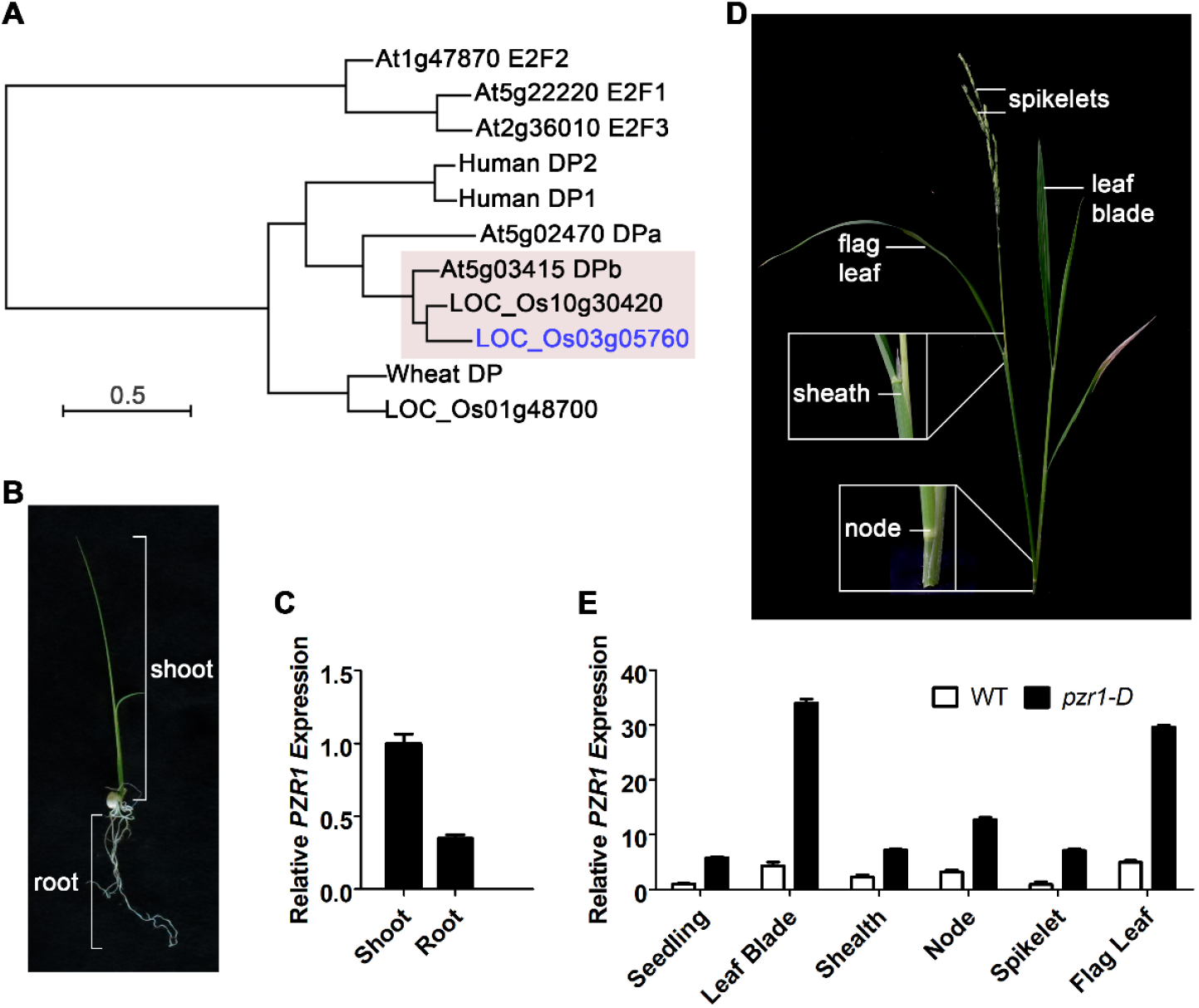
Phylogenetic and expression analysis of PZR1. (**A**) Phylogenetic tree constructed using DP protein sequences from human (UniProt protein ID Q14186 and Q14188), wheat (Q9FET1), Arabidopsis DPa and DPb (Q9FNY2 and Q9FNY3), and three putative rice homologs (Q84VA0, Q84VF4 and Q84VD5). In the pink arabidopsis DPb and two rice homologs, with PZR1 in blue fonts. (**B**) Morphology of 7-day-old rice seedlings showing shoot and root area. (**C**) *PZR1* expression in the shoot and root of wild-type seedlings were examined by RT-qPCR analysis. The bars correspond to standard deviation from three biological replicates. (**D**) Morphology of adult rice plant showing the tissues examined including leaf blades, flag leaf, spikelets, sheath, and node. (**E**) RT-qPCR comparing the expression of *PZR1* in 7-day-old seedlings and different tissues from adult plants.

To elucidate the possible molecular basis of the relationship between BR and cell cycle processes through PZR1, we performed genome-wide transcriptome analysis comparing wild-type and mutant *pzr1-D* seedlings. To avoid detecting individual differences among plants, we sampled three to four seedlings per treatment (one biological replicate) and used two biological replicates per analysis. In total, and considering the consistency of all samples, were identified 1,141 differentially expressed genes (DEGs) among wild-type and mutant seedlings. Of these, 678 genes were differentially expressed only under dark conditions, 229 were differentially expressed in seedlings grown in the light, and 234 were differentially expressed independently of light treatment (Figure 7A). We then identified DEGs that were up- or downregulated in the mutant compared to the wild type and listed the top 20 most significant DEGs under each condition (Figure 7B; Table S1 and S2). To identify characteristics shared by these genes, we performed gene ontology (GO) analysis. Among the most abundant GO terms was cellular processes (GO:0009987), containing 202 DEGs. In the cellular processes category, GO terms such as cell cycle (GO:0007049), cellular component movement (GO:0006928), chromosome segregation (GO:0007059), and cytokinesis (GO:0000910) were enriched. Other enriched terms include cellular component organization (GO:0016043) and cellular component organization or biogenesis (GO:0071840), which are primarily involved in the cell cycle process and regulation (Figure 7C to 7E). In fact, several known rice cell cycle genes were differentially expressed in *pzr1-D* compared to the wild type (Table S3).

**Figure 7.**
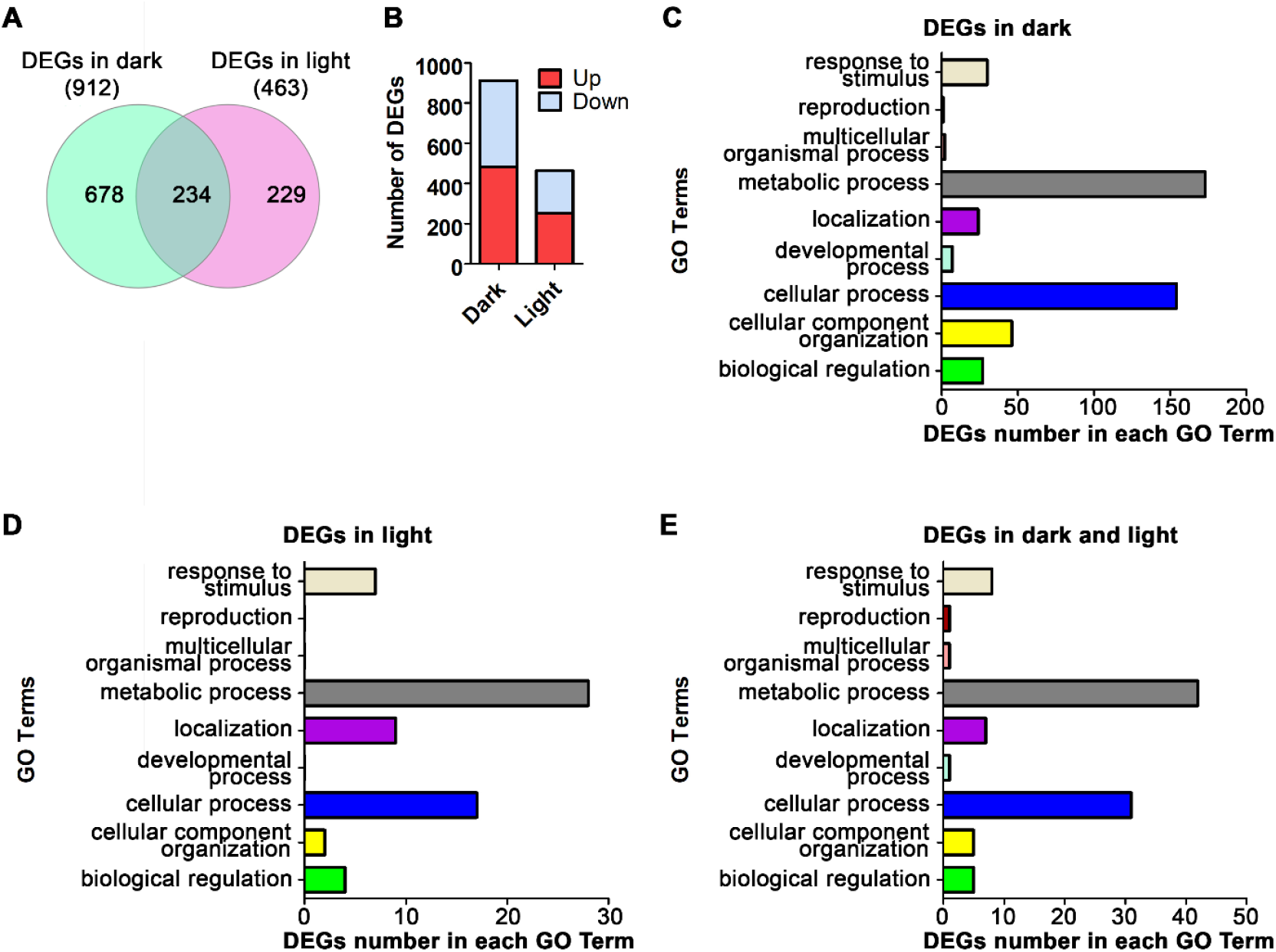
Differentially expressed genes (DEGs) and enriched GO terms for DEGs in mutant and wild-type seedlings. (**A**) Venn diagram of DEGs between the wild type and mutant. A total of 678 DEGs between the wild type and mutant were found in darkness, 229 DEGs were found in the light, and 234 DEGs were found under both conditions. (**B**) Of the 678 DEGs in the dark, 481 were upregulated and 431 were downregulated. In the light, 252 were upregulated and 211 were downregulated. (**C**) List of GO terms for DEGs between wild-type and mutant seedlings maintained under dark or (**D)** light conditions. (**E**) List of GO terms for DEGs under both dark and light conditions.

Finally, we examined the promoter sequences of the top DEGs listed in Tables S1 and S2 and found that 77 out of 80 contained E2F and E2F10SPCNA consensus promoter binding sequences, which match the consensus sequence defined as TTTC[CG]CGC (Vandepoele *et al*., 2005) (Table S4). Together, these results suggest that the *pzr1-D* mutation affects the expression of the genes involved in cell division.

## DISCUSSION

The relative lack of information about BR processes in monocot plants has stimulated the development of new genetic tools and studies. To date, several genes controlling rice architecture and yield were found to be related to BR responses, and many mutants have been identified with potential uses for agronomic improvement (Sakamoto *et al*., 2006, Wu *et al*., 2008, Yang and Hwa, 2008, Wu *et al*., 2016). Like other BR-related mutants in rice, the *pzr1-D* mutant displays semi-dwarfism and an increased number of tillers, two traits that are usually related; dwarf plants generally have more tillers than the wild type. However, unlike other BR mutants, the dwarfism in *pzr1-D* was not accompanied by a significant reduction in seed size. On the contrary, seed size is almost unaffected in this mutant, while an increased tiller number results in an increased number of panicles in the plant, all contributing to improved yield per plant.

In this study, we determined that the mutant phenotype of *pzr1-D* resulted from the activation of *PZR1*, a homolog of an Arabidopsis DP transcription factor gene (Figure 5 and 6). DP is a dimerization partner of the transcription factor E2F that activates the transcription of genes involved in progression to the S phase. We found that overexpression of *PZR1* in rice recapitulated the phenotypes and yield increases to those of the T-DNA activation-tagging lines (Figure 2 and 5). The function of the E2F/DP heterodimer is well conserved in animals and plants, and phylogenetic analysis showed that the heterodimerization domain is well conserved in PZR1, suggesting that PZR1 is likely involved in cell cycle regulation in rice along with E2F, like their Arabidopsis homologs (Figure 6) (De Veylder *et al*., 2002, del Pozo *et al*., 2006). In addition to the results of phylogenetic analysis, other results of this study indicate that PZR1 functions in cell cycle regulation. First, the number cells in roots and leaves were increased in the mutant compared to the wild type, whereas the size of the cells were decreased possibly due to accelerated cell division before cell growth (Figure 4A to 4D; Figure S4). Second, overall shape of the calli derived from *pzr1-D* seeds were smaller and the calli accompanied root-like structures, indicating that the balance between cell division and cell differentiation was clearly altered in the mutant background (Figure 4E and 4F). Additionally, genome-wide transcriptome analysis revealed an association of PZR1 with a number of genes involved in cell cycle regulation (Table S4). Many GO categories related to cellular processes were enriched in the DEGs between wild type and *pzr1-D* (Figure 7). Interestingly, most DEGs have consensus E2F/DP binding sequences in their promoters, again confirming the notion that PZR1 functions like an Arabidopsis DP transcription factor (Figure S6; Table S4).

The dwarf phenotype observed in BR-deficient or -insensitive mutants is mainly caused by decreased cellular elongation, (Kauschmann *et al*., 1996, Szekeres *et al*., 1996), but cell proliferation is altered in this type of mutant as well (Hu *et al*., 2000, Gonzalez-Garcia *et al*., 2011, Zhiponova *et al*., 2013). In Arabidopsis, the overexpression of *E2F* alone produced seedlings with enlarged cotyledons, but the overexpression of *E2F* and its partner *DP* together caused severe dwarfism (De Veylder *et al*., 2002). Similar to our observations in the roots of the *pzr1-D* mutant, cotyledon and roots of *E2F-* and *DP-*overexpressing lines contained more but smaller cells than those of the wild type, and these cells possessed an enlarged nucleus possibly due to enhanced endoreduplication. The extra cells in the *E2F/DP* transgenic plants were proposedly resulted from a prolonged proliferative phase of the cell cycle, which would delay cell differentiation (De Veylder *et al*., 2002). On the other hand, the characteristic phenotype of *pzr1-D* callus may result from shortening of the cell cycle and precocious differentiation (Figure 4E). The PZR1 transcription factor likely serves as an important decision maker determining whether cells should divide.

Interestingly, the overexpression of *DP* alone did not cause any altered phenotype in Arabidopsis seedlings, and overexpression of *E2F* alone resulted in shorter roots (De Veylder *et* al., 2002, Ramirez-Parra *et al*., 2004). However, contrary to both of these findings, overexpression of the rice version of *PZR1*, in Arabidopsis produced seedlings with longer primary roots, increased root hair density, and overall higher root biomass (Figure S5). The phenotypes produced by overexpression of *PZR1* in rice and Arabidopsis, and the observation that this phenotype is opposite to that described for transgenic plants overexpressing *E2F*, prompted us to propose that rice PZR1 and Arabidopsis DP might not have completely overlapping functions and that PZR1 might play a different role aside from merely being a dimer partner of E2F (Figure 5; Figure S5).

The phosphorelay signal transduction pathway plays important roles in BR signaling pathways involving BRI1, BAK1, BSKs, BIN2, and BZR1 (He *et al*., 2002, Li *et al*., 2002, Nam and Li, 2002, Wang *et al*., 2002, Kim and Wang, 2010, Tang *et al*., 2011, Wang *et al*., 2011). Similarly, phosphorylation plays essential roles in cell-cycle control, where cyclin-dependent kinases (CDKs) and cyclins form complexes that phosphorylate targets through progression of cell cycle phases (Inze and De Veylder, 2006). The observation that PZR1 is a cell cycle regulator with BIN2 kinase target motifs in its sequence (Figure S7) led us to hypothesize that it might be a target of phosphorylation by a component of the BR cascade, suggesting a possible link between both processes. One of the most important components of the BR signaling pathway is the protein kinase BIN2, or OsGSK2 in rice, which acts at different levels and even mediates different pathways (Li *et al*., 2001, He *et al*., 2002, Koh *et al*., 2007, Kim *et al*., 2012, Tong *et al*., 2012, Khan *et al*., 2013). A recent example of BIN2-mediated cell cycle regulation was described in rice, where BIN2 was shown to interact and phosphorylate the U-type cyclin, CYC U4 (Sun *et al*., 2015). Most known BIN2 and other GSK3 substrates contain a short consensus sequence, S/TxxxS/T, where S/T corresponds to serine or threonine and x represents any other residue (Zhao *et al*., 2002). Indeed, the sequence of PZR1 harbored many typical motifs (e.g. T^35^xxxS^39^ and S^77^T^78^xxS^81^), raising the possibility that the regulation of PZR1 involves OsGSK2 (Figure S7).

The DP pathway could be manipulated to direct cell division in plants to allow yield and architecture to be adjusted. Since E2F and DP are well-conserved proteins, DP homologs are likely to exist in other cereals of agronomic importance as well. Therefore, similar approaches can be applied to other species through modulating the expression of these homologous genes to boost yield by increasing tiller and panicle number, or to increase drought resistance by amplifying both primary and lateral root production.

## EXPERIMENTAL PROCEDURES

### Plant materials

The wild-type and mutant *Oryza sativa* (rice) plants used in this study were in the Dongjin background. The *pzr1-D* mutant and the other lines examined were identified in a screen of a previously described library of activation-tagging lines (Jeong *et al*., 2002). Rice plants were grown in the paddy field of Seoul National University Agriculture Field in Suwon, Korea (2014 to 2016) or in a greenhouse. For plants in the field, 3-week-old seedlings were transplanted in late May and harvested in mid to late October.

Wild-type Col-0 and transgenic Arabidopsis lines were surface sterilized before sowing on 0.8% agar-solidified medium containing 0.5x Murashige and Skoog (MS) salts and 1% sucrose. After 48 h of stratification at 4°C in darkness, the plates were transferred to a growth room and the plants were grown at 22°C under a 16 h light/8 h dark photoperiod in white light (80 µmol m^− 2^ s ^−1^). If needed, the seedlings were transferred to soil after 10 days of growth in MS medium.

### Chemical treatments and morphological analysis

Rice seeds were sterilized for 30 min with 50% sodium hypochlorite and washed 4–5 times with water prior to planting. Before each treatment, the seeds were germinated on filter paper soaked with water for 2 days. Newly germinated seeds at similar growth stage were subjected to treatments at a planting depth of 1 cm. For Pcz treatment, the seeds were planted in coarse vermiculite soil soaked with water supplemented with the indicated Pcz concentrations. Pcz (100 mM) dissolved in DMSO was used as a working solution, and DMSO alone was added to water as a control or mock treatment. Plants were maintained in total darkness or in the light (80-100 µmol m^− 2^ s ^−1^ intensity) under long-day conditions as indicated in each experiment. After 7 days of treatment, images were taken, and growth parameters were measured from the images using ImageJ software. Overall plant height was measured from the end of the root to the highest leaf, whereas the length of the main root was used to determine root length. Seed length and area were calculated from the digital photographs using ImageJ software. All statistical analyses were performed using GraphPad Prism 5 software. Significance was evaluated using Student’s *t*-test.

### Genomic DNA and genotyping

DNA extraction was performed using a DNA Prep Kit (BioFACT) following the manufacturer’s recommendations. The DNA was quantified using a spectrophotometer system (BioTek) controlled with the Gen5 Data Analysis software interface. Genotyping was performed using 50 ng DNA template in two sets of PCR. In one set, specific primers for *LOC_Os03g05760* and the surrounding region were used to amplify the wild-type allele. The other set was performed using a primer specific to the left border of the T-DNA (pGA2715 LB) to amplify the mutant allele in which the T-DNA was inserted. This analysis allowed us to identify homozygous and heterozygous mutant plants for each line, as well as segregating wild-type plants. A line with only the wild-type allele was considered to be segregating wild type (w/w), and a line with only the mutant allele amplified was considered to be homozygous (T/T). Lines in which both alleles were amplified were considered to be heterozygous for the insertion (w/T). The sequences of the primers used for PCR are listed in Table S5.

### Lamina-joint bending assay

The lamina bending bioassay was performed as described (Wada *et al*., 1984, Zhang *et al*., 2012), with slight modifications. Seeds were sterilized and germinated on filter papers and transferred to 0.5x MS medium, followed by incubation for 7 days in darkness. Segments containing the second-leaf lamina joint were cut from uniformly growing seedlings. The segments were floated on distilled water for 24 h in darkness to remove any chemical residues from the plant that might alter the experiment, and the samples were checked to ensure that all lamina angles were similar prior to treatment. Uniform samples with similar lamina angles were floated on distilled water containing the indicated concentration of brassinolide (BL). All procedures, including sectioning of the samples and transfer to various solutions, were performed in a dark room to avoid exposure to light as much as possible. The segments were incubated for 48 h in darkness under different treatments and photographed. The photographs were used to measure the angle between the lamina and the blade sheath using ImageJ software. All statistical analysis was performed using GraphPad Prism 5 software, and significance was evaluated using Student’s *t*-test.

### RNA isolation and gene expression analysis

The total RNA used for RT-qPCR analysis was isolated from rice or Arabidopsis tissues using an RNeasy system (Qiagen) following the manufacturer’s instructions. The cDNA was synthesized from 2 µg RNA using M-MLV reverse transcriptase (ELPIS). RT-qPCR analysis was performed on an Applied Biosystems StepOne Real-Time PCR System with Power SYBR Green PCR Master Mix as previously described (Corvalan and Choe, 2017) using the primers listed in Table S5. For the Arabidopsis OX lines, gene expression data in RT-qPCR were normalized against OX line 8, which showed low transcript levels in the RT-PCR analysis, as *Loc_Os03g05760* is a rice gene and is therefore not present in Col-0 control plants.

### Phylogenetic analysis

Protein sequences showing similarity to PZR1 were retrieved using the BLAST server on the Phytozome website (https://phytozome.jgi.doe.gov/pz/portal.html). Sequences of Arabidopsis, rice, wheat, and human proteins were obtained from the UniProt server (http://www.uniprot.org), and phylogenetic analysis was performed using Clustal Omega (http://www.ebi.ac.uk/Tools/msa/clustalo/) and BoxShade software (http://embnet.vital-it.ch/software/BOX_form.html) to identify and visualize conserved sequences. The resulting phylogenetic tree was generated using the Neighbor-joining method (Clustal Omega) and modified using FigTree v1.4.3. A list of the protein sequences used as input can be found in Table S6.

### Cloning and plant transformation

Vectors used to produce rice and Arabidopsis plants overexpressing rice *OsDPB/PZR1* under the control of the 35S promoter were constructed as follows: RNA was extracted, and cDNA was synthesized from 7-day-old rice seedlings with specific primers (Table S5) to amplify the full-length CDS of interest. The resulting products (696 bp) corresponding to the gene of interest was purified and cloned into the entry vector pENTR/SD/D-TOPO (Invitrogen), followed by cloning into the destination vector pEarleyGate101(C-YFP-HA), which is compatible with the Gateway system. For Arabidopsis transformation, *Agrobacterium* strain GV3101 was transformed with the vectors, and the LBA4404 strain was used for rice. The constructs were transformed into plants using conventional *Agrobacterium*-mediated techniques, and transgenic seedlings were selected on MS medium supplemented with 20 mg/L BASTA. BASTA-resistant and -sensitive plants were identified, and a chi-square test was carried out to test monogenic segregation pattern.

### Callus induction

Rice seeds were sterilized and placed on filter paper until complete dry. The seeds (12–18) were germinated and cultured on 2N6 medium plates containing Duchefa’s Chu (N6) powder (4 g L^- 1^), sucrose (30 g L^-1^), L-proline (2.9 g L^-1^), casein (0.3 g L^-1^), myo-inositol (0.1 g L^-1^), 2.4-D (2 mg L^-1^), and Phytagel (4 g L^-1^) at pH 5.8. The plates were incubated in a growth chamber under dark conditions at 32°C for the indicated number of days before being weighed and photographed for analysis with ImageJ software.

### Microscopy

Root images were obtained under a Leica TCS SP9 confocal laser-scanning microscope. Root samples were excised from 7-day-old seedlings 1 cm above the root tip and submerged in 10 μg/mL propidium iodide (PI) solution for 3 minutes. The samples were rinsed twice in distilled water prior to observation. Images were compiled and analyzed using LAS X software version 3.0.2. For cell counting and size determination, cells inside a 60 μm^2^ square drawn in the meristematic zone 300 μm above the root tip were examined. Leaves were photographed under a Primo Vert inverted microscope (Zeiss), and ImageJ software was used for measurement. Each leaf of a 7-day-old seedling was dissected transversally down the middle, and images were taken and used to compare genotypes.

### Transcriptome analysis and identification of DEGs between wild type and *pzr1-D* mutant seedlings

Total RNA was extracted from seedlings using Trizol (Sigma-Aldrich) reagent following the manufacturer’s instructions. Treatments were performed in triplicate for seedlings of two genotypes, wild type (WT) and *pzr1-D* (MUT), which were grown in the light or total darkness. A total of 12 samples were prepared and subjected to quality inspection. Of these, duplicates of each treatment with the best purity value were used to construct sequencing libraries. Quality inspection, library preparation, and raw data processing were conducted by a contracting service (Theragen Etex, http://www.theragenetex.com/). Quality assessment was performed on an Agilent Bioanalyzer 2100 with RNA integrity number (RIN) >7 (>9 for most samples). Libraries were generated using an Ultra™ RNA Library Prep Kit for Illumina and sequenced on an Illumina HiSeq 4000 system. Differentially expressed genes (DEGs) were identified in the samples as wild-type seedlings grown in darkness versus *pzr1-D* seedlings grown in darkness (WT dark vs. *pzr1-D* dark) and wild-type seedlings grown in light versus *pzr1-D* seedlings grown in light (WT light vs. *pzr1-D* light). The expression level of each gene was calculated using HTseq software and normalized. DEGs were identified based on log_2_ (fold changes) ≥ 1 and a corrected P-value (Q-value) of ≤ 0.05. Gene ontology (GO) analysis, Venn diagram construction, and promoter analysis were performed using tools available online at http://pantherdb.org/, http://www.interactivenn.net/, and http://plantpan2.itps.ncku.edu.tw/gene_group.php?#multipromoters, respectively.

## AUTHOR CONTRIBUTIONS

S.C. designed the research and analyzed the data; S.Y. performed experiments and analyzed the data; G.A. provided the T-DNA activation tagging mutant population and C.C. performed the research and wrote the paper with input from the other authors.

## ACKNOWLEDGEMENTS

We are grateful to SI Kwon, SM Chu, and EH Kang for technical assistance. This research was supported, in part, by grants from the Cooperative Research Program for Agricultural Science and Technology Development (Project No. PJ01168501), the Rural Development Administration, and the National Research Foundation of Korea (NRF; grant No. 2015R1A2A1A10051668).

## SUPPORTING INFORMATION

### Supplementary Figures

**Supplemental Figure 1.**
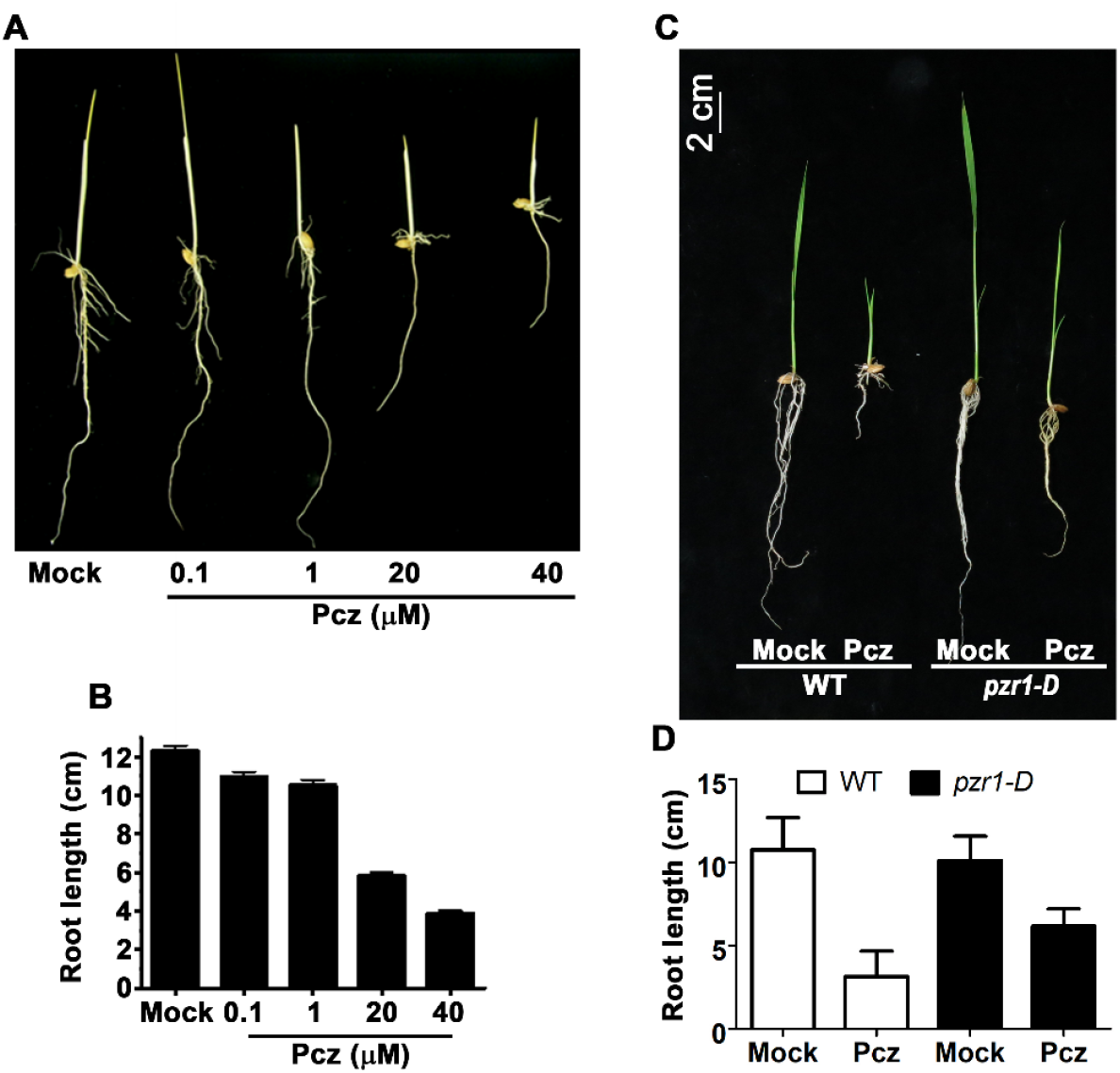
Dose response and light studies of propiconazole effect in rice. (**A**) Morphology and (**B)** measurement of root lengths of plants after 10 days of treatment with 0 (Mock) to 40 μM of the BR inhibitor Pcz. (**C**) Morphology and (**D)** measurement of root lengths of seedlings after 10 days of treatment with 30 μM Pcz under normal light conditions. In the graphs, error bars represent standard deviation of 10 or more samples per treatment.

**Supplemental Figure 2.**
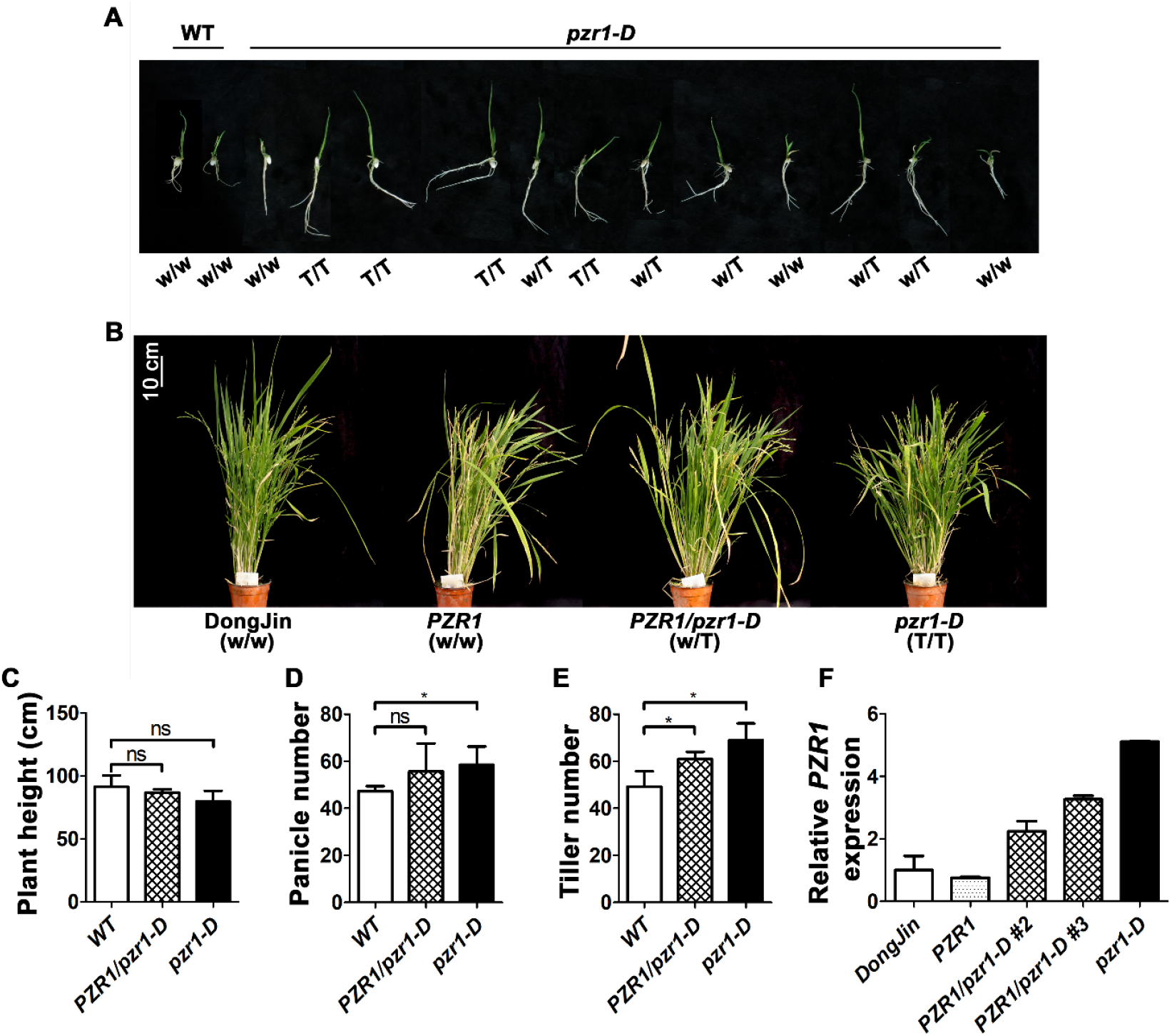
Propiconazole sensitivity and phenotypes of *pzr1-D* progeny. (**A**) Morphology of 7-day-old seedlings grown in Pcz-supplemented medium. The different genotypes are represented as w/w for the wild type, T/T as homozygous mutant (where T represents T-DNA), and w/T for the heterozygote. (**B**) A representative plant of each genotype is shown; wild type Dongjin (w/w), segregating wild type or *PZR1* (w/w), heterozygous *PZR1/pzr1*-D (w/T), and homozygous mutant *pzr1-D* (T/T). Graphs comparing (**C**) plant height, (**D**) panicle number, and (**E**) tiller number. (**F**) RT-qPCR analysis of *PZR1* expression in each genotype.

**Supplemental Figure 3.**
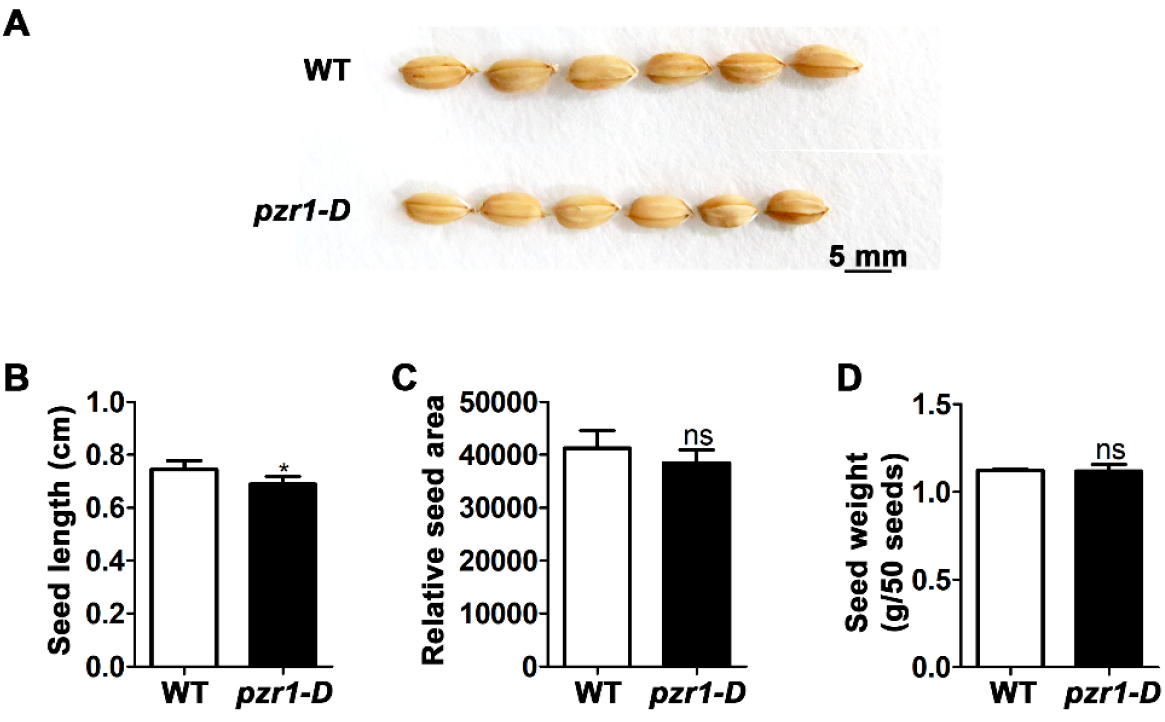
Analysis of *pzr1-D* seeds. (**A**) Wild-type and mutant seeds were compared in terms of (**B**) seed length, (**C**) relative area, and (**D**) weight. A total of 200 seeds per genotype were measured, and the mean was used to plot the graphs. Error bars represent standard deviation, and significant differences were determined using Student’s *t*-test. *, P<0.05; ns, non-significant difference.

**Supplemental Figure 4.**
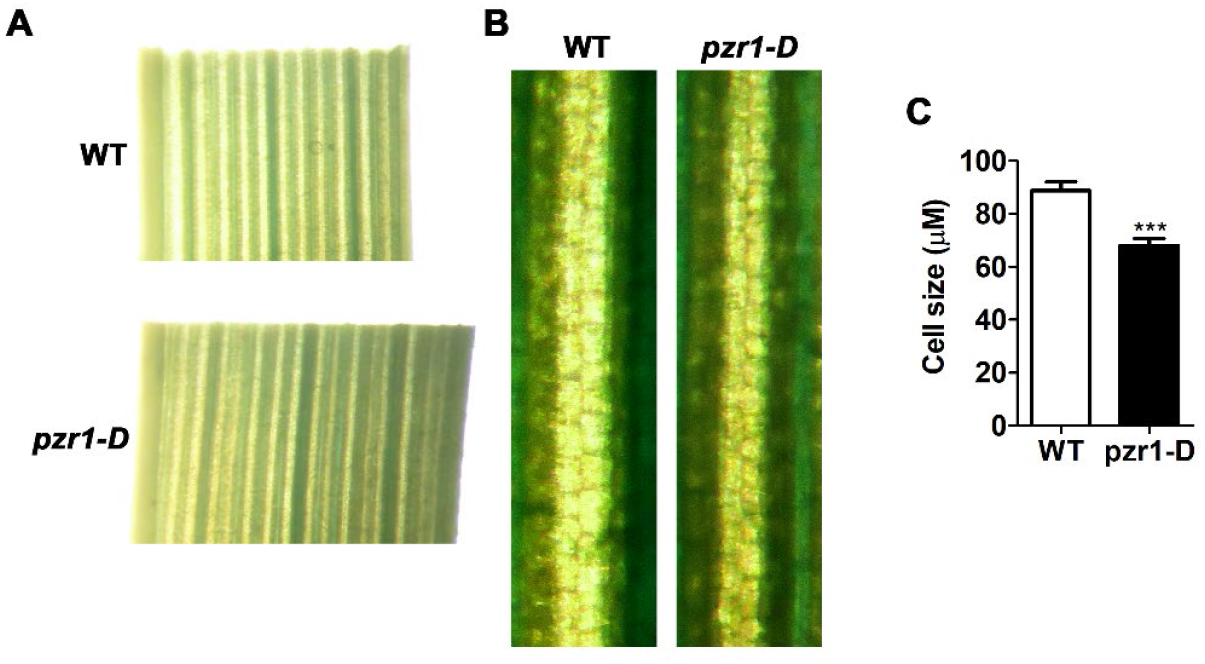
Microscopy analysis of leaves from wild type and *pzr1-D* mutant seedlings. (**A**) Images of leaves from 7-day-old seedlings wild-type and mutant seedlings dissected transversally down the middle observed under 10X magnification. (**B**) Images of samples observed under 20X magnification and (**C**) used to measure cell size. Six slides with samples per genotype were analyzed, each with 20 to 30 cells measured to obtain the average and standard deviation shown in the graph. Significance was determined using Student’s *t*-test. ***, P<0.0001.

**Supplemental Figure 5.**
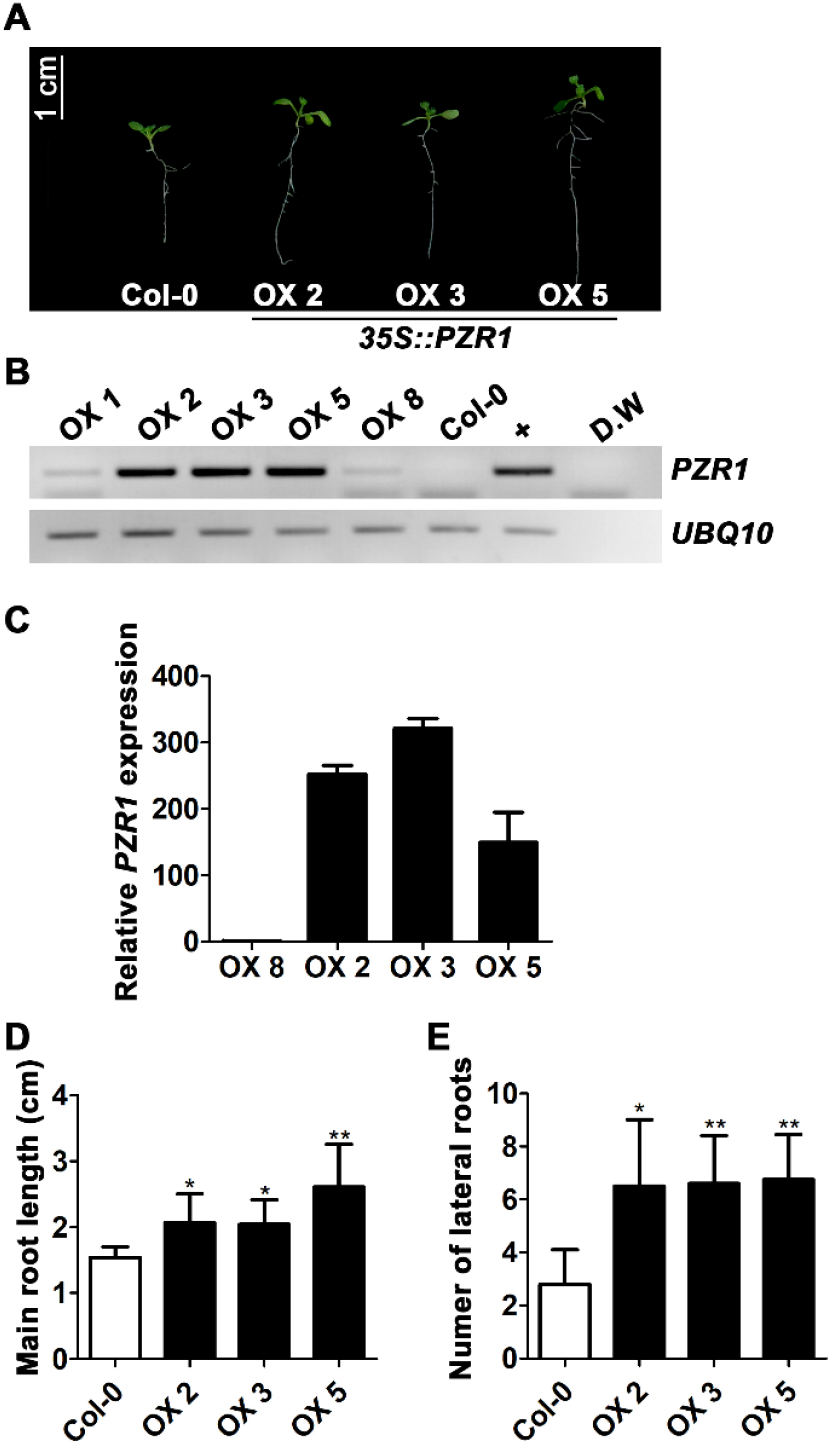
Phenotypes and gene expression levels of Arabidopsis plants heterologously expressing rice *PZR1*. (**A**) Morphology of non-transformed Col-0 wild-type plant and representative seedlings from three independent transgenic lines. (**B**) RNA samples were collected from seedlings of independent transgenic lines to measure the expression levels of rice *PZR1*. Positive and negative controls are represented with the sign + and D.W (distilled water) respectively. Overexpression lines OX 1 and OX 8, which produced very low levels of the fragment corresponding to *PZR1*, were used as a reference. Arabidopsis *UBIQUITIN 10* was used as an internal control. (**C**) RT-qPCR analysis of *PZR1* expression in transgenic plants using OX 8 as a reference, which expresses *PZR1* at very low levels. (**D**) Primary root length and (**E**) number of lateral roots are shown as the average value (n=10). The error bars represent standard deviation, and significant differences were determined using Student’s *t*-test. *, P<0.05; **, P<0.001; and ***, P<0.0001.

**Supplemental Figure 6.**
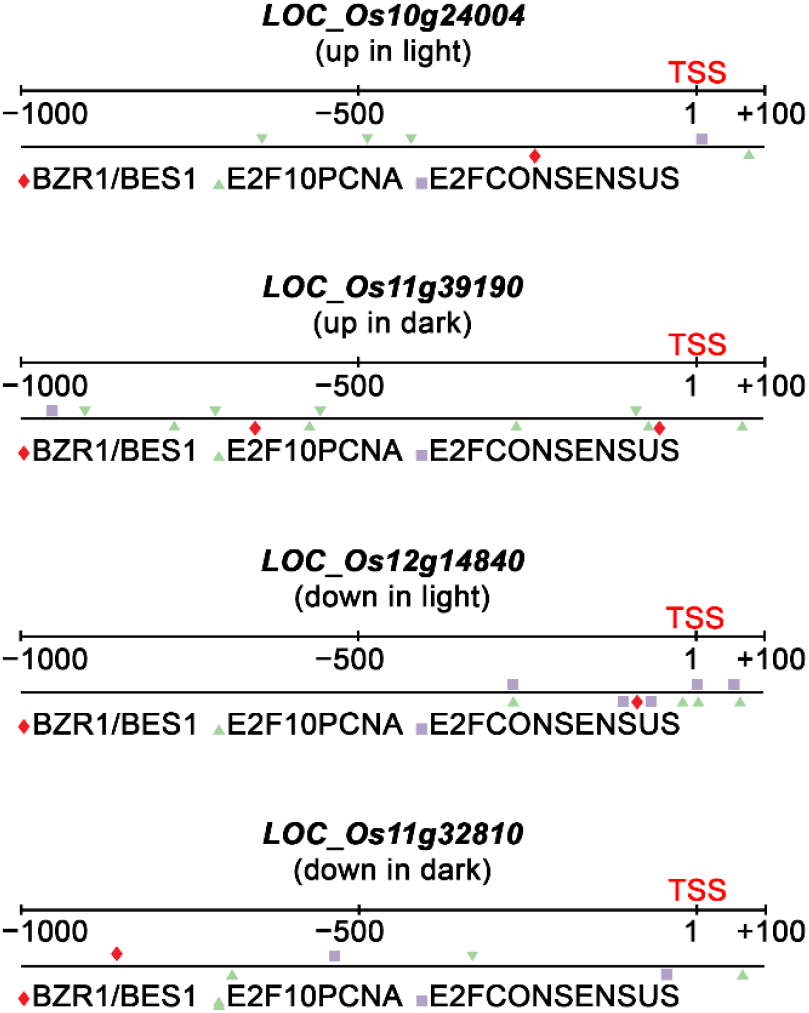
Promoter regions of the top DEGs. Schematic representation of four different gene promoter regions and the locations of E2F/DP and BZR1/BES1 consensus cis-acting elements. TSS defines the transcription start site, red rhombus indicates position of BZR1/BES1 sites, green triangle E2F10 PCNA and purple square the E2F consensus sites.

**Supplemental Figure 7.**
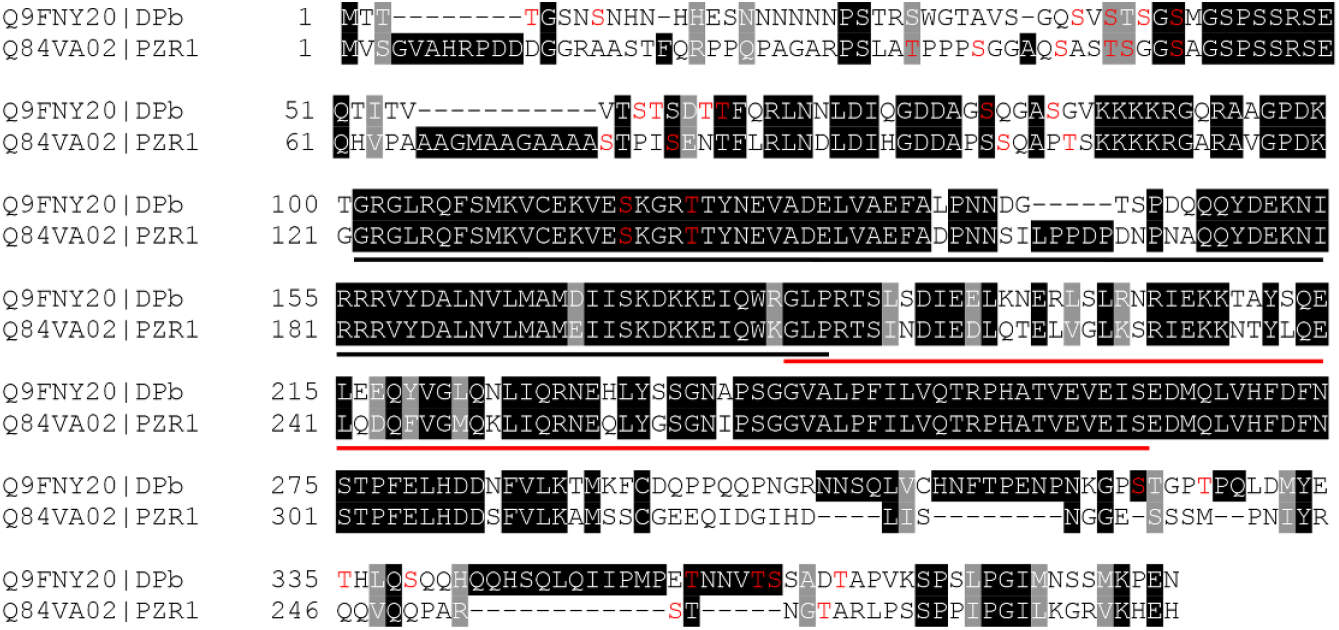
Multiple Sequence Alignment of DP proteins. Multiple sequence analysis of Arabidopsis DPb and rice homolog PZR1 proteins. In DPb, the black underline delimits DNA binding domain (amino acids 101-184) while the red line represents the heterodimerization domain (182-263). The red letters S and T represent Serine and Threonine that follows the S/TxxxS/T pattern of phosphorylation by BIN2 and homologs.

### Supplementary Tables

**Supplemental Table 1.**
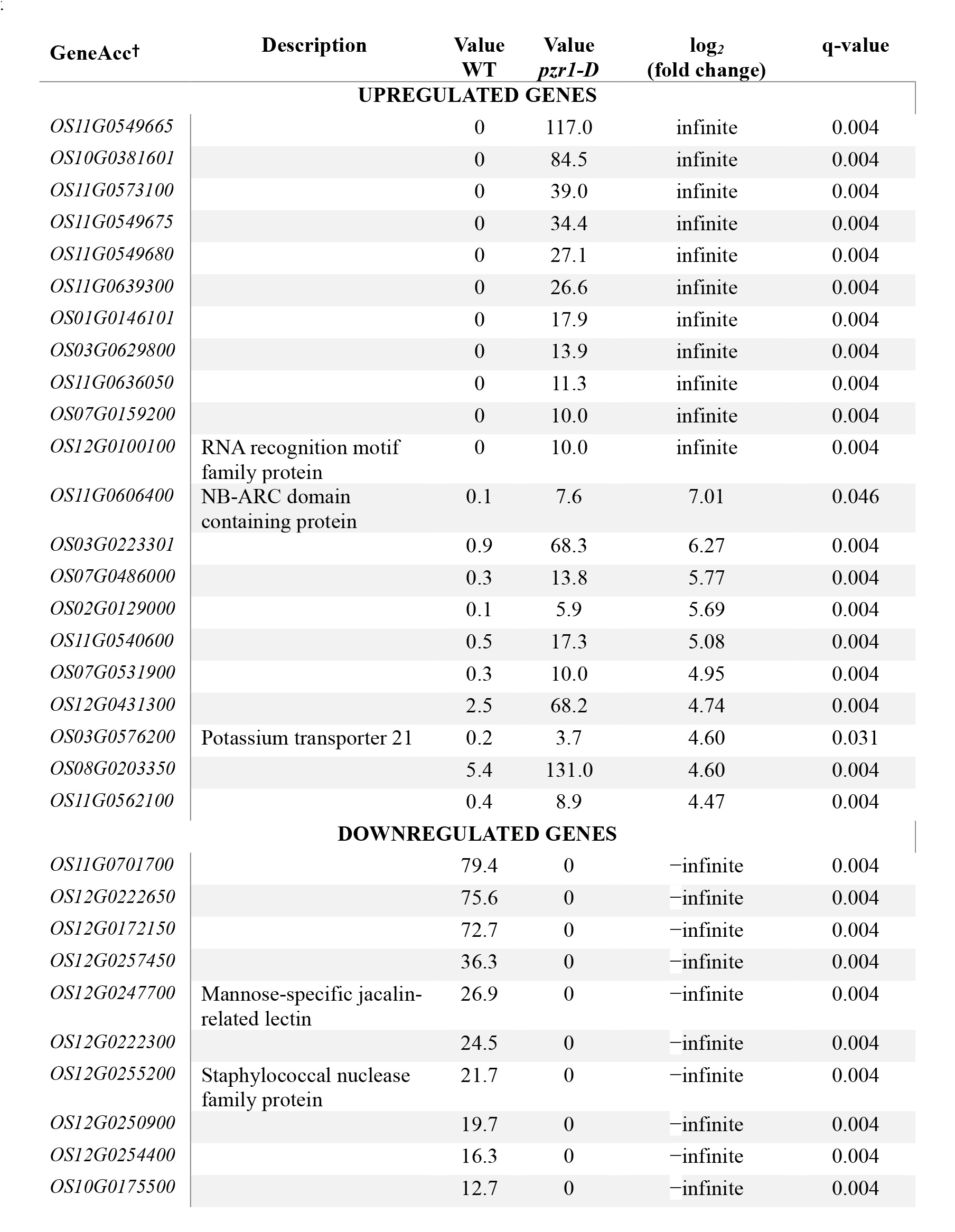
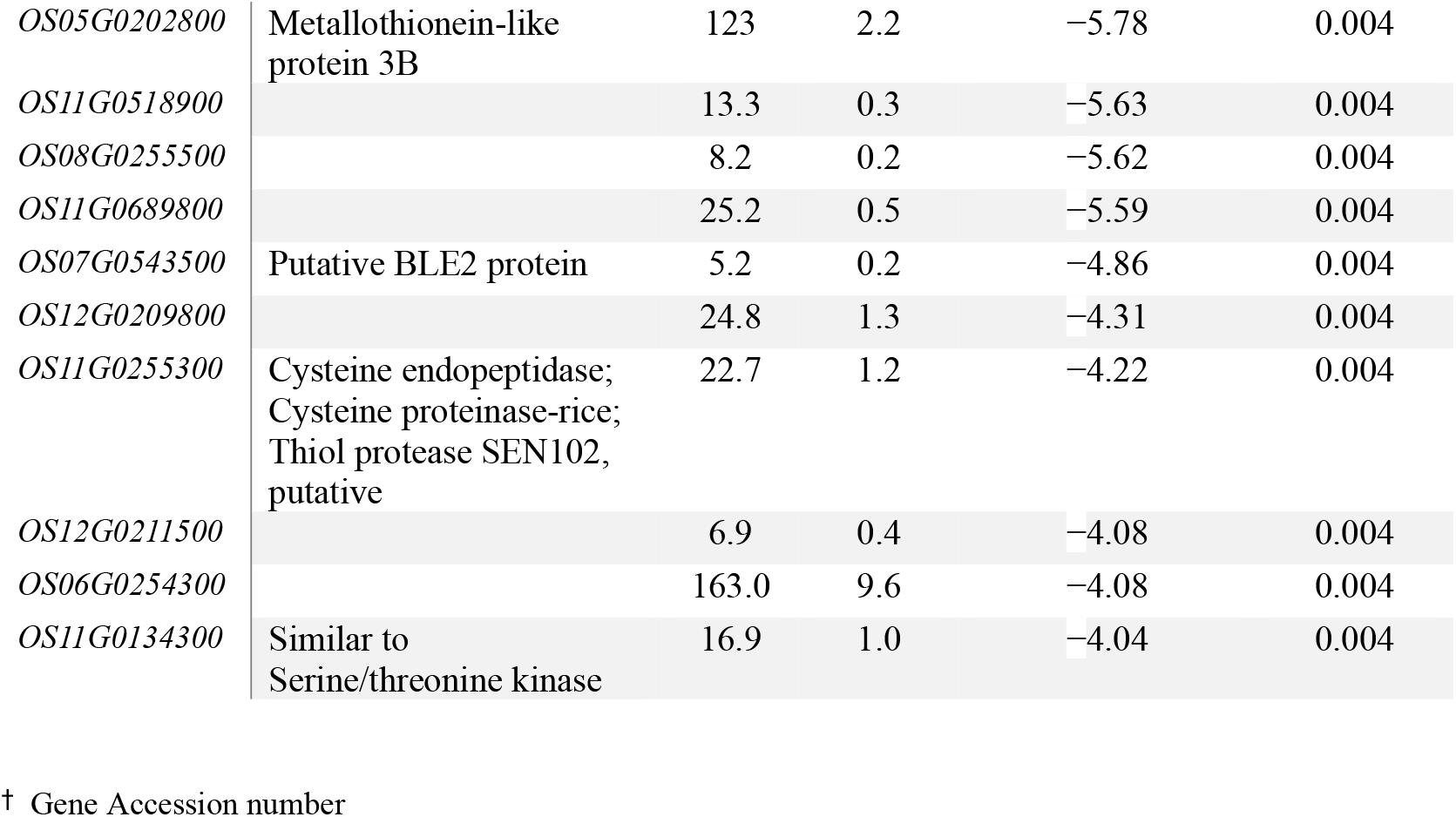
List of the top 20 most significant differentially expressed genes (DEGs) in *pzr1-D* compared with the wild type (WT) in the light.

**Supplemental Table 2.** List of the top 20 most significant differentially expressed genes (DEGs) in *pzr1-D* compared with the wild type (WT) in darkness.

| GeneAcc† | Description | Value<br>WT | Value<br>*pzr1-D* | log<sub>2</sub><br>(fold change) | q-value |
| --- | --- | --- | --- | --- | --- |
| **UPREGULATED GENES** |  |  |  |  |  |
| *OS09G0467700* |  | 0 | 32.6 | infinite | 0.003 |
| *OS01G0146101* |  | 0 | 20.9 | infinite | 0.003 |
| *OS01G0148100* |  | 0 | 13.8 | infinite | 0.003 |
| *OS07G0187001* |  | 0 | 12.0 | infinite | 0.003 |
| *OS11G0640300* | Leucine Rich Repeat<br>family protein,<br>expressed | 0 | 8.5 | infinite | 0.003 |
| *OS12G0257400* |  | 0 | 7.5 | infinite | 0.003 |
| *OS07G0159200* |  | 0 | 7.4 | infinite | 0.003 |
| *OS01G0845950* |  | 0 | 7.0 | infinite | 0.003 |
| *OS11G0691100* |  | 0 | 5.9 | infinite | 0.003 |
| *OS07G0153150* |  | 0.2 | 68.4 | 8.16 | 0.003 |
| *OS11G0605100* | NB-ARC domain<br>containing protein | 0.1 | 11.8 | 7.18 | 0.006 |
| *OS03G0223301* |  | 0.5 | 68.2 | 7.08 | 0.003 |
| *OS08G0367300* |  | 0.1 | 7.1 | 6.67 | 0.019 |
| *OS11G0618700* |  | 0.1 | 9.6 | 6.44 | 0.005 |
| *OS07G0162450* |  | 0.5 | 44.8 | 6.37 | 0.003 |
| *OS02G0129000* |  | 0.1 | 8.9 | 6.30 | 0.003 |
| *OS03G0299700* |  | 1.6 | 123 | 6.25 | 0.003 |
| *OS11G0549680* |  | 0.4 | 26.1 | 6.00 | 0.003 |
| *OS07G0486000* |  | 0.2 | 15.8 | 5.97 | 0.003 |
| *OS11G0569800* | Receptor kinase,<br>putative, expressed | 0.2 | 11.2 | 5.83 | 0.003 |
| *OS09G0467700* | - | 0 | 32.6 | infinite | 0.003 |
| **DOWNREGULATED GENES** |  |  |  |  |  |
| *OS12G0250900* | - | 20.9 | 0.5 | -5.33 | 0.003 |
| *OS12G0406000* | - | 6.4 | 0.2 | -5.14 | 0.003 |
| *OS11G0696600* | - | 26.1 | 0.8 | -5.09 | 0.003 |
| *OS11G0532600* | - | 5.5 | 0.2 | -5.01 | 0.010 |
| *OS01G0520180* | - | 81.5 | 2.6 | -4.99 | 0.003 |
| *OS07G0297400* | - | 93.3 | 2.9 | -4.97 | 0.003 |
| *OS07G0535200* | - | 6.74 | 0.2 | -4.78 | 0.003 |
| *OS12G0425800* | - | 56.2 | 2.1 | -4.74 | 0.003 |
| *OS11G0685200* | - | 7.5 | 0.3 | -4.62 | 0.003 |
| *OS12G0204600* | NB-ARC domain<br>containing protein | 6.5 | 0.3 | -4.62 | 0.003 |
| *OS12G0239300* | - | 281.0 | 12.1 | -4.53 | 0.003 |
| *OS07G0677100* | Peroxidase | 40.3 | 1.8 | -4.50 | 0.003 |
| *OS07G0103000* | - | 15.1 | 0.7 | -4.49 | 0.010 |
| *OS05G0414400* | - | 6.2 | 0.3 | -4.32 | 0.003 |
| *OS11G0693800* | - | 18.2 | 0.9 | -4.25 | 0.003 |
| *OS05G0369900* | - | 8.1 | 0.5 | -4.09 | 0.003 |
| *OS11G0687100* | Von Willebrand factor<br>type A domain<br>containing protein,<br>expressed | 41.0 | 2.5 | -4.04 | 0.003 |
| *OS12G0425500* | - | 3.7 | 0.2 | -4.04 | 0.003 |
| *OS08G0255500* | - | 6.6 | 0.4 | -3.99 | 0.003 |
| *OS11G0689800* | - | 16.1 | 1.2 | -3.77 | 0.003 |

† Gene Accession number

**Supplemental Table 3.** List of differentially expressed rice cell cycle genes in *pzr1-D* compared with the wild type (WT).

| Common name | Description | Sample | Value in WT | Value in *pzr1-D* | log<sub>2</sub> (fold change) |
| --- | --- | --- | --- | --- | --- |
| *OsRAA1* | Flowering-promoting factor 1-like protein 1 | WT vs *pzr1-D* in the dark | 6.625 | 19.519 | 1.559 |
|  |  | WT vs *pzr1-D* in the light | 5.698 | 16.287 | 1.515 |
| *CycB1;1* | Cyclin-B1-1 | WT vs *pzr1-D* in the dark | 65.441 | 39.659 | -0.723 |
|  |  | WT vs *pzr1-D* in the light | 21.389 | 22.576 | 0.078 |
| *Psd1* | Similar to Lipid transfer protein. | WT vs *pzr1-D* in the dark | 649.943 | 336.993 | -0.948 |
|  |  | WT vs *pzr1-D* in the light | 192.136 | 278.211 | 0.534 |
| *CDKB;1* | Similar to Cyclin-dependent kinase B1-1 | WT vs *pzr1-D* in the dark | 25.206 | 15.539 | -0.698 |
|  |  | WT vs *pzr1-D* in the light | 13.253 | 11.259 | -0.235 |
| *CycT1;3* | Cyclin-dependent kinase A-2 | WT vs *pzr1-D* in the dark | 18.959 | 12.393 | -0.613 |
|  |  | WT vs *pzr1-D* in the light | 9.255 | 8.796 | -0.073 |
| *CDKG;1* | Cyclin-dependent kinase G-1 | WT vs *pzr1-D* in the dark | 18.775 | 32.877 | 0.808 |
|  |  | WT vs *pzr1-D* in the light | 39.671 | 40.669 | 0.036 |
| *PCNA* | Proliferating cell nuclear antigen | WT vs *pzr1-D* in the dark | 129.495 | 68.486 | -0.919 |
|  |  | WT vs *pzr1-D* in the light | 37.115 | 45.642 | 0.298 |
| *CKS1* | Cyclin-dependent kinases regulatory subunit 1 | WT vs *pzr1-D* in the dark | 49.949 | 31.799 | -0.651 |
|  |  | WT vs *pzr1-D* in the light | 36.218 | 35.047 | -0.047 |
| *CDKG;2* | Cyclin-dependent kinase G-2 | WT vs *pzr1-D* in the dark | 17.678 | 24.627 | 0.478 |
|  |  | WT vs *pzr1-D* in the light | 28.401 | 24.079 | -0.238 |
| *CycB2;1* | Cyclin-B2-1 | WT vs *pzr1-D* in the dark | 24.121 | 15.741 | -0.616 |
|  |  | WT vs *pzr1-D* in the light | 8.909 | 8.037 | -0.149 |
| *CycU1;1* | Cyclin-P1-1 | WT vs *pzr1-D* in the dark | 0.497 | 1.333 | 1.424 |
|  |  | WT vs *pzr1-D* in the light | 4.273 | 8.237 | 0.947 |
| *CycD2;2* | Similar to Cyclin-D3-1 | WT vs p zr1-D in the dark | 10.231 | 6.013 | -0.767 |
|  |  | WT vs p zr1-D in the light | 5.588 | 5.224 | -0.097 |
| *CycB2;2* | Cyclin-B2-2 | WT vs p zr1-D in the dark | 46.600 | 27.177 | -0.778 |
|  |  | WT vs p zr1-D in the light | 13.102 | 14.476 | 0.144 |
| *CycD6;1* | Cyclin-D6-1 | WT vs p zr1-D in the dark | 1.046 | 0.290 | -1.849 |
|  |  | WT vs p zr1-D in the light | 0.257 | 0.373 | 0.537 |
| *CycD2;1* | Cyclin-D2-2 | WT vs p zr1-D in the dark | 17.749 | 9.653 | -0.879 |
|  |  | WT vs p zr1-D in the light | 6.290 | 6.813 | 0.115 |
| *CDKB2;1* | Cyclin-dependent kinase B2-1 | WT vs p zr1-D in the dark | 58.090 | 34.057 | -0.770 |
|  |  | WT vs p zr1-D in the light | 24.909 | 26.196 | 0.073 |
| *CDKB;2* | Cyclin-dependent kinase B2-1 | WT vs p zr1-D in the dark | 58.090 | 34.057 | -0.770 |
|  |  | WT vs p zr1-D in the light | 24.909 | 26.196 | 0.073 |
| *CycA3;2* | Cyclin-A3-2 | WT vs p zr1-D in the dark | 9.151 | 5.852 | -0.645 |
|  |  | WT vs p zr1-D in the light | 3.704 | 4.441 | 0.262 |

**Supplemental Table 4.**
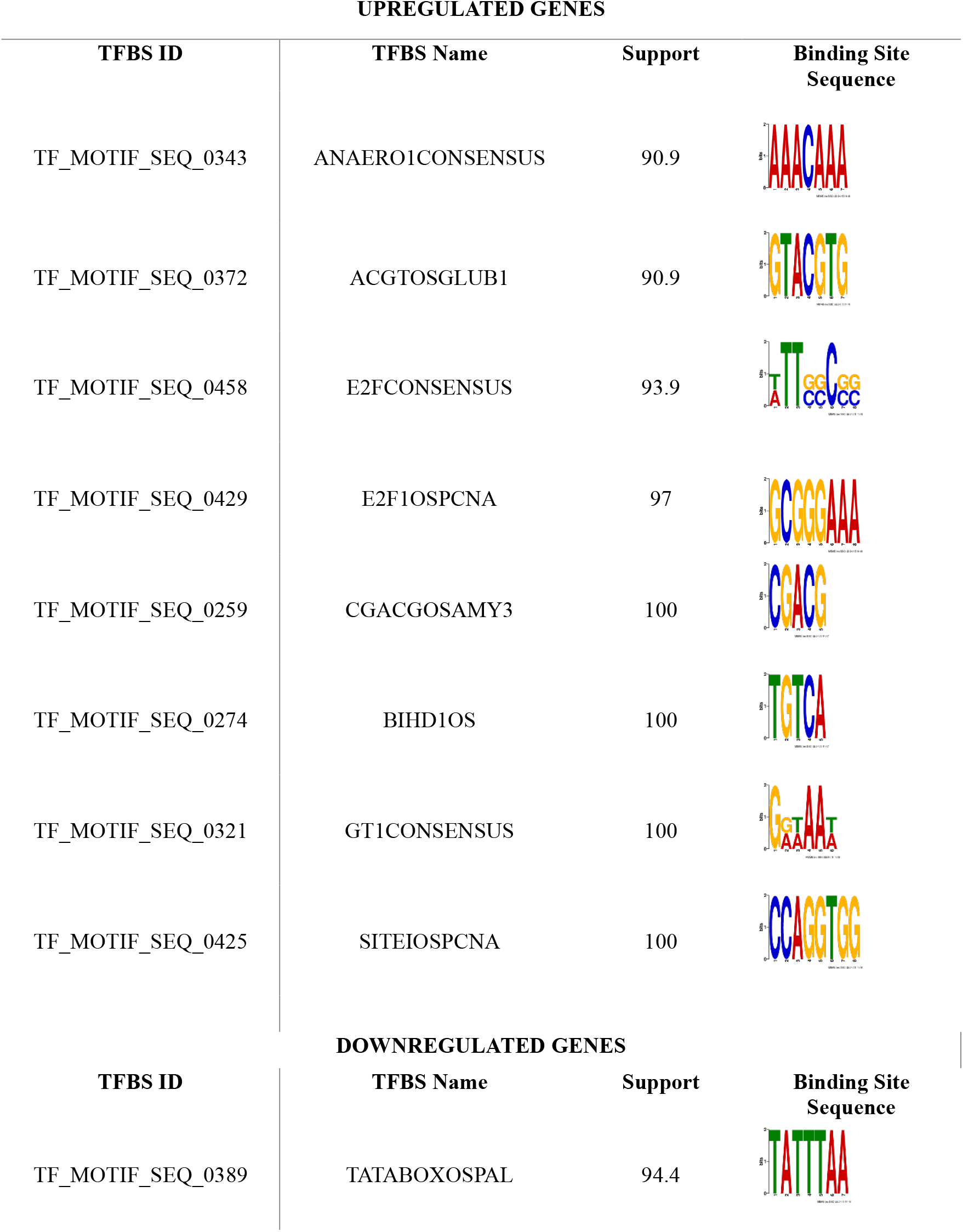
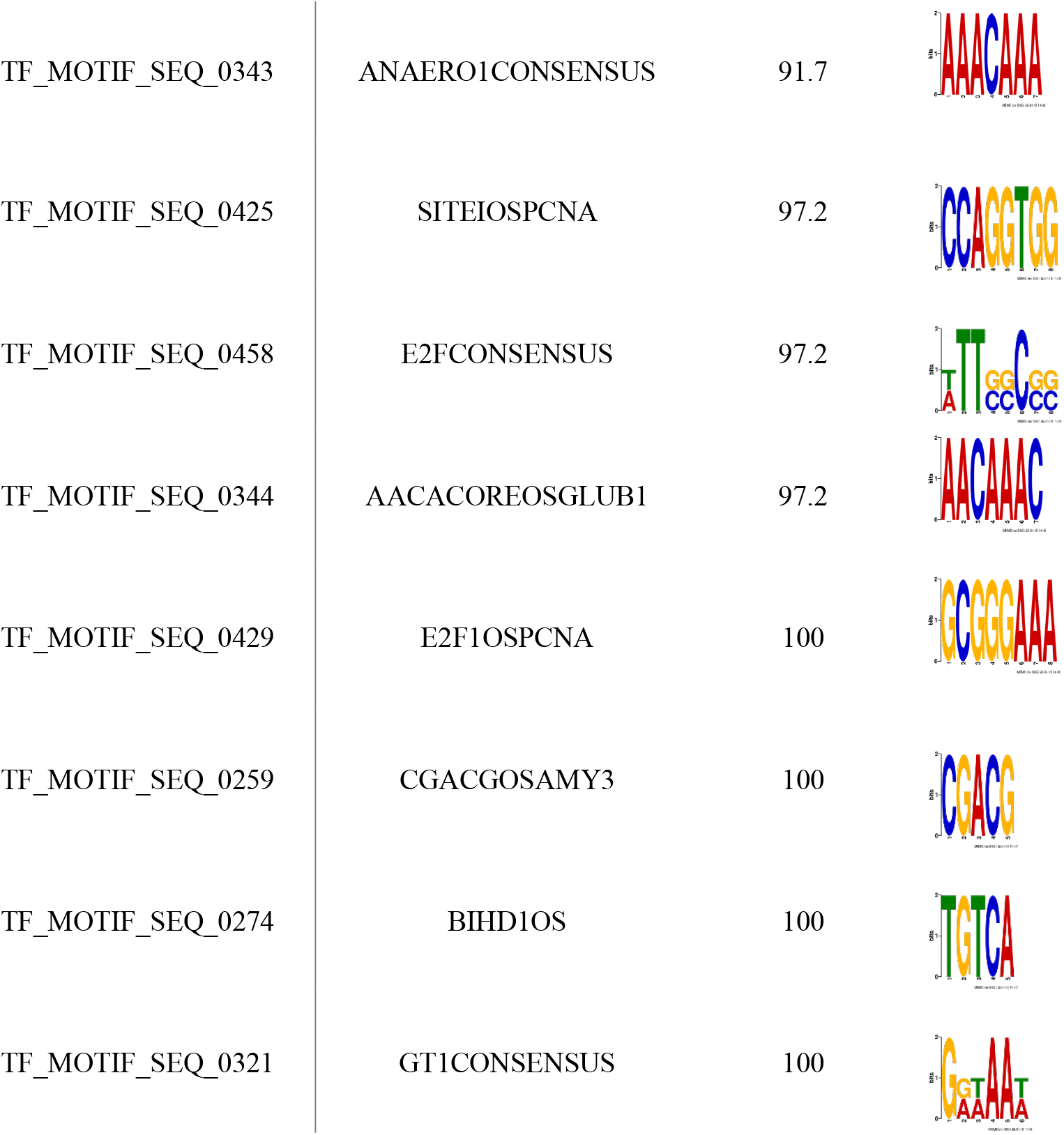
Transcription Factor Binding Sites (TFBS) in the promoters of the top 20 most significant differentially expressed genes (DEGs) from each condition.

| UPREGULATED GENES |  |  |  |
| --- | --- | --- | --- |
| TFBS ID | TFBS Name | Support | Binding Site Sequence |
| TF_MOTIF_SEQ_0343 | ANAERO1CONSENSUS | 90.9 |  |
| TF_MOTIF_SEQ_0372 | ACGTOSGLUB1 | 90.9 |  |
| TF_MOTIF_SEQ_0458 | E2FCONSENSUS | 93.9 |  |
| TF_MOTIF_SEQ_0429 | E2F1OSPCNA | 97 |  |
| TF_MOTIF_SEQ_0259 | CGACGOSAMY3 | 100 |  |
| TF_MOTIF_SEQ_0274 | BIHD1OS | 100 |  |
| TF_MOTIF_SEQ_0321 | GT1CONSENSUS | 100 |  |
| TF_MOTIF_SEQ_0425 | SITEIOSPCNA | 100 |  |
| DOWNREGULATED GENES |  |  |  |
| TFBS ID | TFBS Name | Support | Binding Site Sequence |
| TF_MOTIF_SEQ_0389 | TATABOXOSPAL | 94.4 |  |
| TF_MOTIF_SEQ_0343 | ANAERO1CONSENSUS | 91.7 |  |
| TF_MOTIF_SEQ_0425 | SITEIOSPCNA | 97.2 |  |
| TF_MOTIF_SEQ_0458 | E2FCONSENSUS | 97.2 |  |
| TF_MOTIF_SEQ_0344 | AACACOREOSGLUB1 | 97.2 |  |
| TF_MOTIF_SEQ_0429 | E2F1OSPCNA | 100 |  |
| TF_MOTIF_SEQ_0259 | CGACGOSAMY3 | 100 |  |
| TF_MOTIF_SEQ_0274 | BIHD1OS | 100 |  |
| TF_MOTIF_SEQ_0321 | GT1CONSENSUS | 100 |  |

**Supplemental Table 5.**
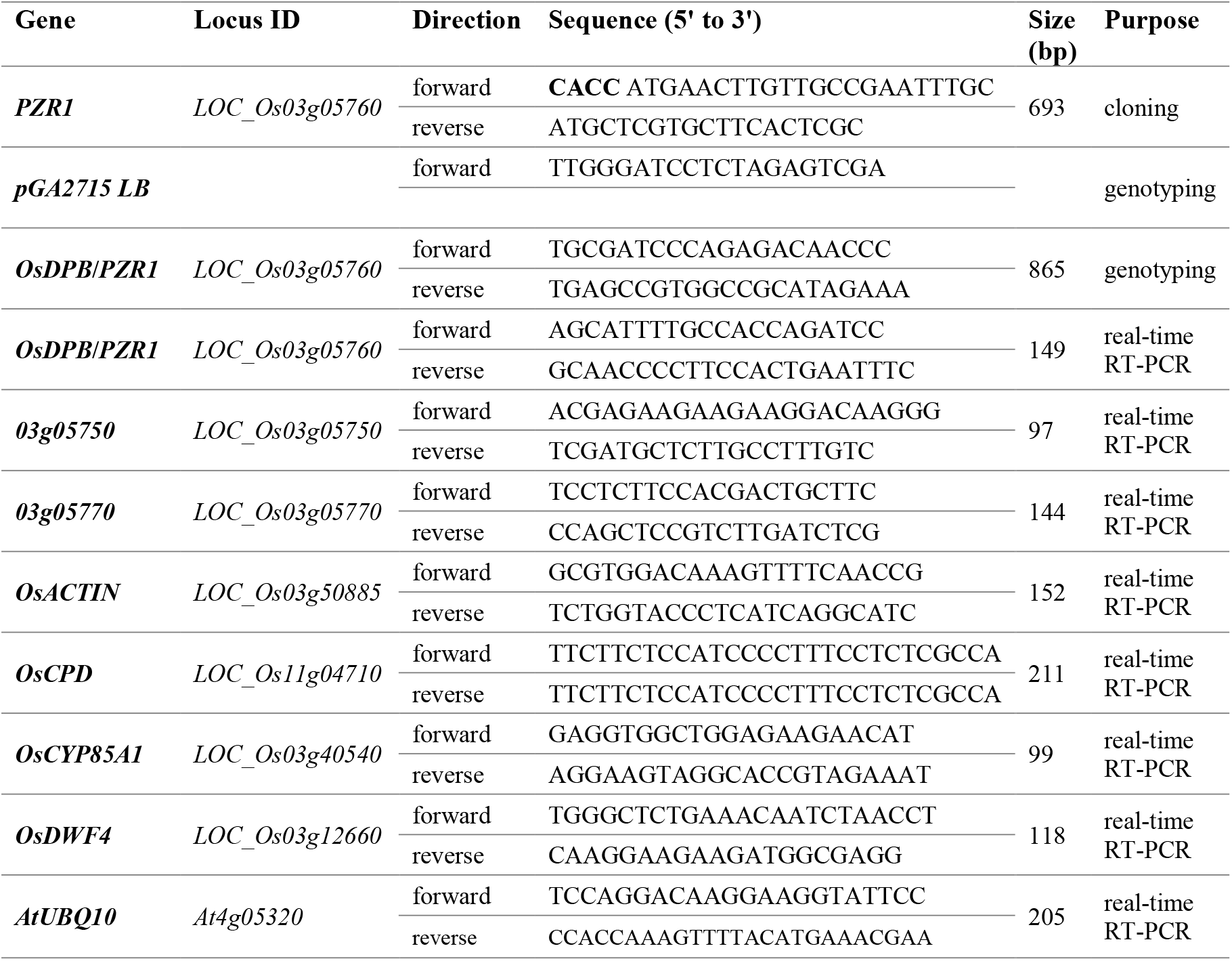
Primers used in this study.

**Supplemental Table 6.**
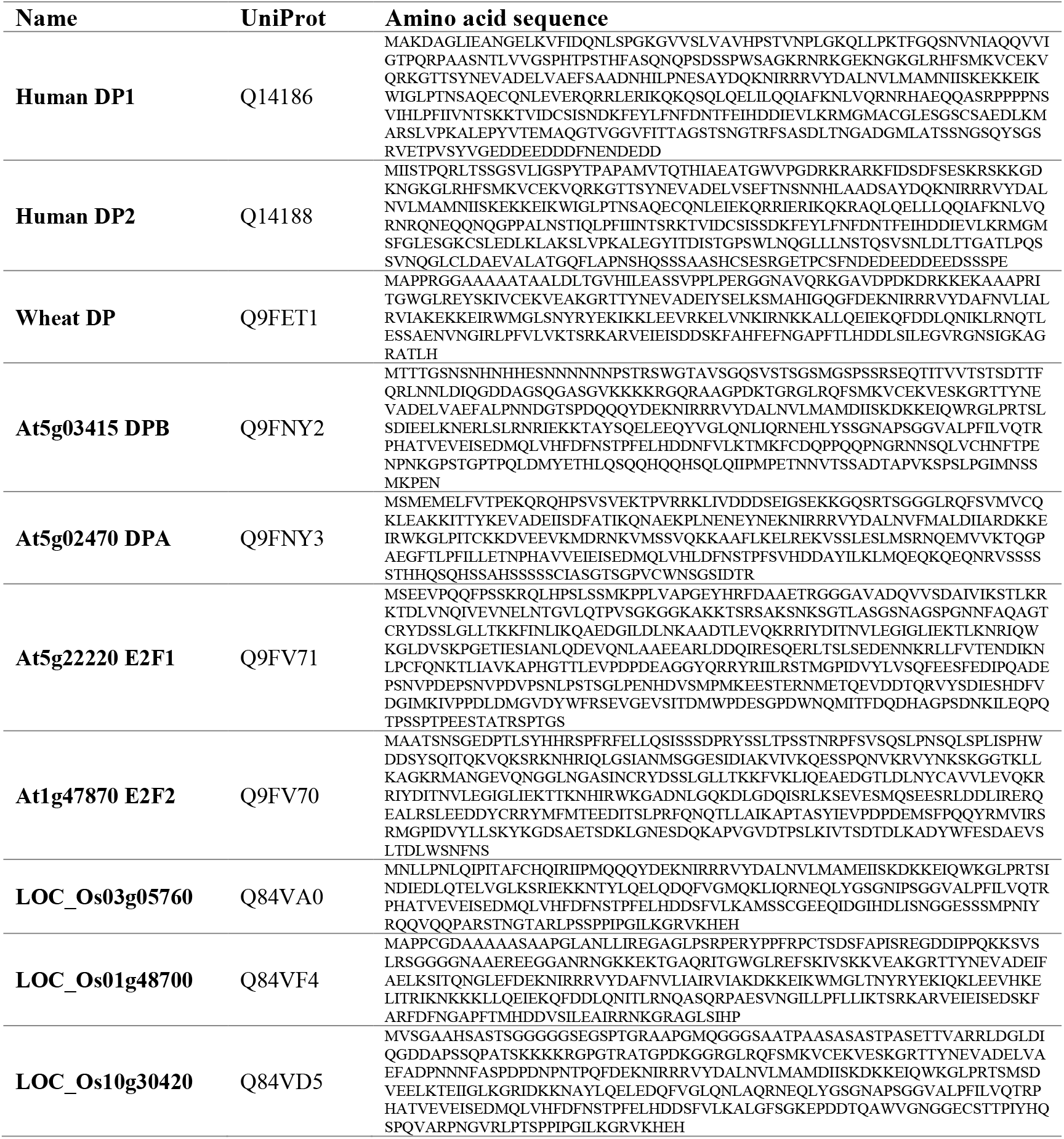
List of amino acid sequences used for phylogenetic analysis.

